# A universal polymer signature in Hi-C resolves cohesin loop density and supports monomeric extrusion

**DOI:** 10.1101/2025.09.04.674214

**Authors:** Kirill Polovnikov, Dmitry Starkov

## Abstract

Cohesin organizes mammalian chromosomes by extruding DNA loops, but whether this process involves single cohesin complexes or paired “dimers” in vivo has remained controversial. Here, we identify a universal physical signature of cohesin loop density in Hi-C contact statistics at short genomic separations. In this regime, contact statistics simplify dramatically and loops can be treated as static, enabling an analytically solvable equilibrium polymer theory for a looped chain with finite contact detection. The theory predicts a characteristic dip in the slope of the contact probability curve that depends only on two parameters: cohesin loop density and the contact capture radius determined by the experimental protocol. Applying this framework to diverse mammalian Hi-C datasets, we infer a conserved loop density of approximately six loops per megabase. Independent cohesin copy-number measurements from quantitative imaging and mass spectrometry match this value, indicating that loop extrusion is predominantly monomeric in vivo, although rare or transient higher-order assemblies cannot be excluded. More broadly, this framework integrates Hi-C, imaging, and biochemical measurements within a single physical model, providing new mechanistic insight into how motor proteins organize chromatin in vivo.

**Significance Statement:** How many cohesin complexes are required to extrude a chromatin loop in living cells remains a central question in genome biology. We develop a polymer-physics framework that links short-range Hi-C features to the density of cohesin-mediated loops in vivo. The theory explains a characteristic contact signature that depends solely on loop density. Applying this readout across mammalian datasets, we infer ∼ six loops per megabase in interphase chromatin, implying that ∼ 60 − 70% of the genome is folded within cohesin-mediated loops. Independent imaging and biochemical measurements yield comparable values, supporting a predominantly monomeric mode of loop extrusion in vivo while not excluding rare or transient dimeric events. By bridging polymer physics with genome-wide contact maps, this framework provides a quantitative route to dissect how motor proteins organize chromosomes.

## Introduction

The three-dimensional organization of chromosomes plays a central role in genomic processes, including gene regulation, replication, repair, and segregation [1]. Mammalian interphase chromosomes exhibit a hierarchical folding pattern: at the megabase scale, active and inactive chromatin segregate into compartments, while sub-megabase structures such as topologically associating domains (TADs) emerge as key organizational units [2, 3]. A leading mechanism shaping this architecture is loop extrusion by cohesin, a ring-shaped SMC complex that reels in DNA to form dynamic loops [4–6].

A central unresolved question concerns the *stoichiometry* of cohesin during loop extrusion in vivo—that is, how many cohesin complexes are required to extrude a single chromatin loop. Although single cohesin complexes can extrude DNA in vitro [7, 8], super-resolution imaging and biochemical studies in cells have reported both monomeric and dimeric assemblies [9– 11]. Existing approaches have been unable to resolve this ambiguity for three main reasons. First, extrusion is highly dynamic, and stalled or clustered cohesins can appear as “dimers” in microscopy even when acting as monomers [11]. Second, direct visualization of interphase loops is technically challenging because loops form and dissolve rapidly. Third, copy-number measurements alone cannot determine the number of cohesins per loop because the key structural parameter—the *density of loops*—remains unknown. Bridging molecular-scale motor activity with genome-wide chromosome architecture therefore requires a quantitative physical framework.

Chromosome conformation capture (Hi-C) provides exactly the type of genome-wide structural information needed to connect molecular mechanisms with chromosome architecture. Hi-C experiments generate a contact map in which each entry reflects how frequently two genomic loci are found in close spatial proximity. By pooling all pairs of loci separated by the same genomic distance *s*, one obtains the contact probability curve *P* (*s*), a robust physical footprint of chromatin folding. Across cell types and species, *P* (*s*) displays several characteristic features [6, 12], which polymer theory has linked to underlying loop architecture. A peak in the slope of *P* (*s*) around *s* ≈ 100 kb reflects the typical loop size [6]. By contrast, loop density has remained elusive, obscured by other structural parameters at large genomic scales. Focusing on short scales, we identify another conserved feature: a local minimum (“dip”) in the *P* (*s*) slope at separations of tens of kilobases (Fig. 1F). Its physical origin and quantitative relationship to cohesin-mediated loop extrusion have remained unclear, partly because Hi-C only detects contacts within a finite effective radius determined by cross-linking and fragment size [13, 14]. Without accounting for this protocol coarse-graining, key structural signatures in *P* (*s*) can be blurred or misinterpreted. Here we introduce a polymer-physics framework that quantitatively links short-range Hi-C contact statistics to the density of cohesin-mediated loops as a single structural parameter. Specifically, the model reduces the short-scale Hi-C signal to just two interpretable quantities: a structural one (loop density) and a protocol-dependent one (capture radius). As we show, both parameters can be inferred robustly from the Hi-C contact map. The analytical framework is validated through active-extrusion simulations and two independent experimental perturbations, one modulating cohesin abundance and the other altering Hi-C resolution. Applying this approach to diverse mammalian datasets reveals a remarkably conserved loop density of ∼ 6 *±* 1 loops per megabase. This Hi-C-based inference matches independent measurements of RAD-21 (cohesin subunit) counts in microscopy and mass-spectrometry, supporting a predominantly monomeric stoichiometry of extruding cohesin in living interphase nuclei.

**Figure 1.**
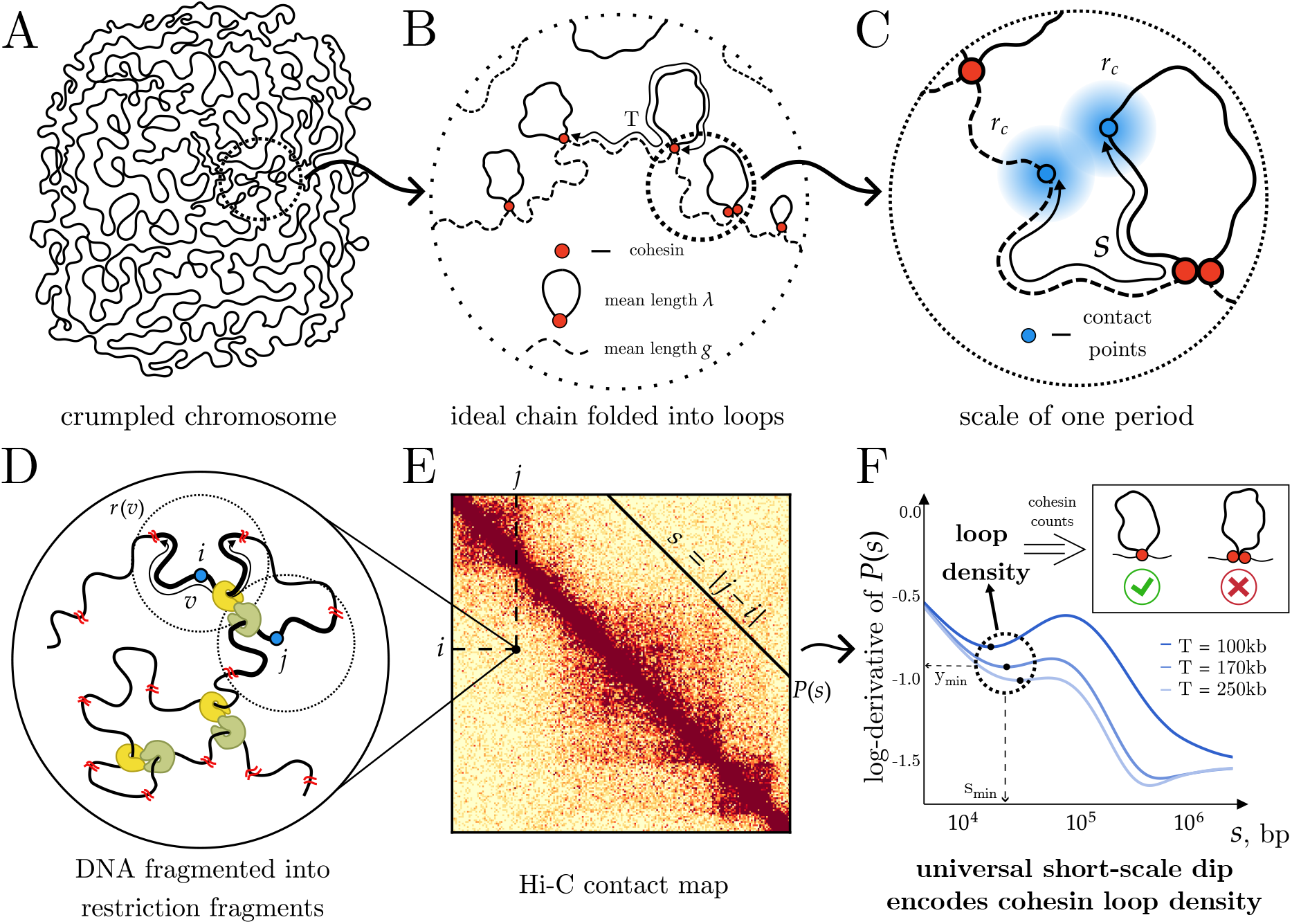
From chromosome territories to a short-scale polymer model of Hi-C contacts.(A) Interphase chromosomes form compact, crumpled polymers within nuclear territories. (B) At intermediate scales, chromatin behaves as an ideal polymer bridged by cohesin (red), forming loops (solid) and gaps (dashed) with mean lengths *λ* and *g*; their sum defines the loop–gap period *T* (loop density *T* ^−1^). **(C)** At short genomic separations (*s < λ, g*), folding is governed primarily by *T*, and contacts are detected only within the Hi-C capture radius *r*_*c*_ (blue halo). **(D)** Cross-linking and restriction digestion generate fragments of length *v* (mean *v*_0_) with spatial extent *r*(*v*). **(E)** Ligation of proximal fragments yields contacts contributing to *P* (*s*). **(F)** Theoretical log-derivative curves of *P* (*s*) for varying *T* (with *λ* = 80 kb fixed) and 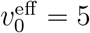 Kb show a universal short-scale dip (*s*_min_, *y*_min_) that fingerprints loop density. Comparing inferred loop densities with cohesin copy-number measurements supports predominantly monomeric extrusion in vivo.

## Results

### A short-scale Hi-C fingerprint reveals cohesin loop density

Chromosome folding is a multiscale phenomenon, with distinct statistical regimes emerging at different genomic distances (Fig. 1A-C). At scales larger than the cohesin loop size (*λ* ≈ 100kb), this organization becomes particularly complex, arising from the interplay of multiple parameters that define the loop architecture, topological entanglements, and the intrinsic activity of loop extrusion. Our previous work examined this complex interplay using an equilibrium approximation of frozen loops on a crumpled polymer, qualitatively reproducing the characteristic shape of Hi-C contact statistics *P* (*s*) [6, 12]. By contrast, as we show here, at scales below the typical loop size chromosome folding simplifies dramatically, enabling robust and quantitative inference of loop density.

Across mammalian Hi-C maps, the log-derivative of the contact probability *P* (*s*)—the slope on a log–log plot—exhibits a remarkably conserved local minimum (“dip”) at genomic separations of a few tens of kilobases (Fig. 1F, Fig. 2A). To explain this universal feature of chromatin folding, in what follows we construct a polymer model that couples a quenched loop-gap structure – with loops of mean size *λ* and gaps of mean size *g*, defining a period *T* = *λ* + *g* (loop density *T* ^−1^) – to the capture radius of the Hi-C protocol, *r*_*c*_, which determines when two loci are experimentally registered as a contact. For convenience, we express this capture radius in equivalent genomic units using an effective fragment length 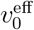 whose spatial size corresponds to *r*_*c*_. The model yields three falsifiable predictions: (i) the dip arises only from the *combined* action of loops and a finite capture radius; (ii) its genomic position scales as 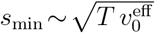; and (iii) its depth *y*_min_ depends only on the ratio 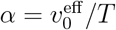. Together, these relations allow the two model parameters to be extracted directly from the dip’s position and depth, revealing the underlying loop density from the short-range Hi-C signal (Fig. 1F).

**Figure 2.**
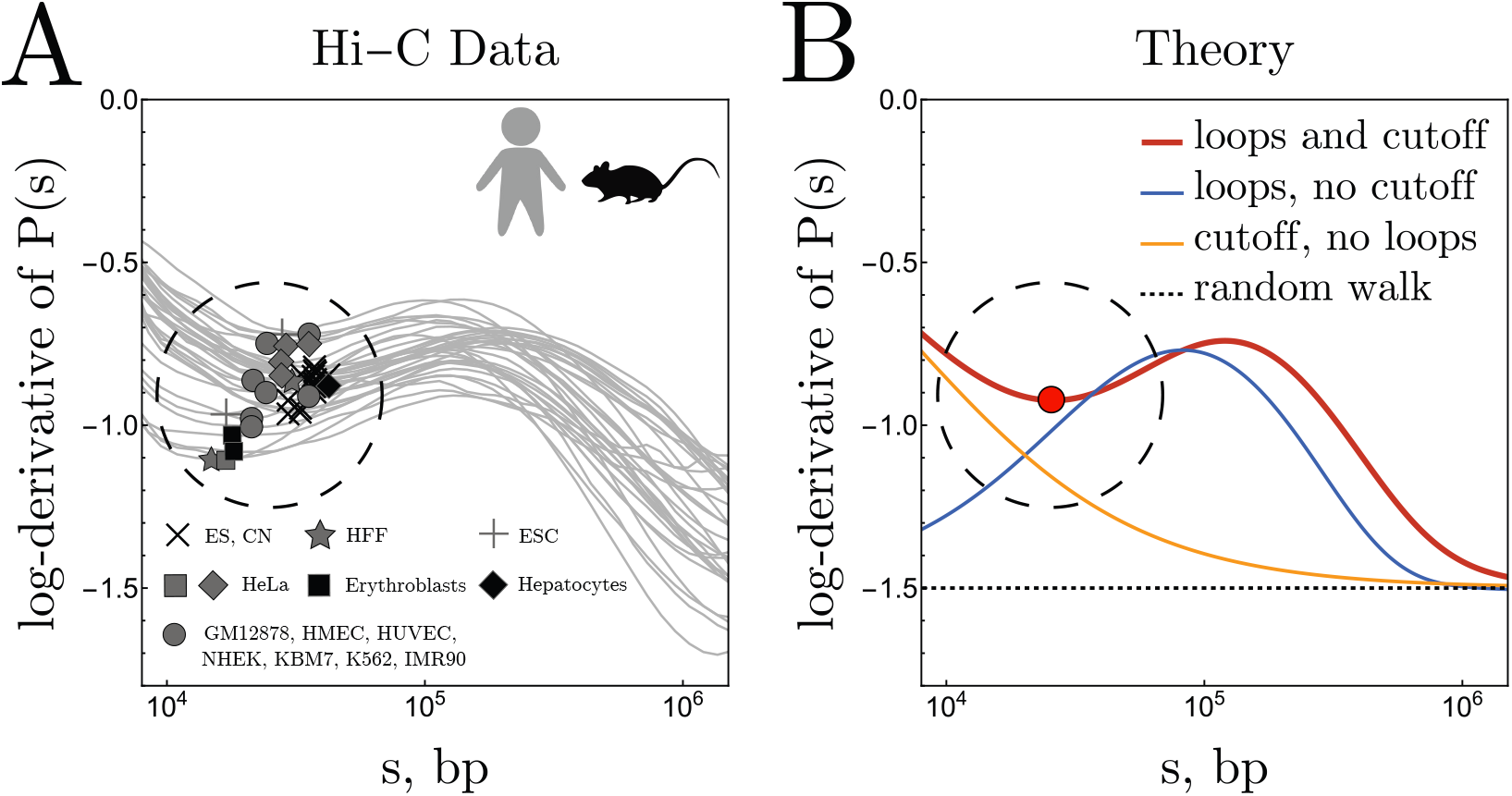
Loops and finite capture radius are both required to produce the universal short-range dip in Hi-C.**(A)** Log-derivative of *P* (*s*) for 33 human and mouse Hi-C datasets, all showing a conserved minimum at tens of kilobases. **(B)** Polymer-model predictions. Only the full model with both loops and finite capture radius (red) reproduces the dip; models lacking loops (orange) or finite capture radius (blue) do not. Parameters: *λ* = 120 kb, *g* = 50 kb (*T* = 170 kb); 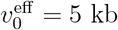. Dashed line: ideal-chain slope −3*/*2.

### Loop-free baseline

To facilitate exposition of our theory, we first analyse chromatin folding in the absence of loops, a state that can be experimentally induced by cohesin depletion or degradation. Within the genomic range relevant to our analysis (*s* ∼ 5–50 kb), loop-free chromatin behaves as an ideal polymer of fractal dimension *d*_*f*_ = 2, satisfying the relation *r*(*s*) = *b s*^1*/*2^. Here 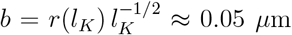 is the effective Kuhn length in spatial units, obtained by scaling the spatial extent of a Kuhn segment (*r*(*l*_*K*_) ≈ 0.1 *µ*m) by its contour length *l*_*K*_ ≈ 4 kb [15]. In this genomic window, chromatin has not yet entered the entangled regime characteristic of larger scales, and stiffness effects remain confined to sub-kilobase separations, making ideal-chain statistics a reasonably accurate baseline.

Although this polymer baseline is relatively simple, the Hi-C detection process remains a significant and non-trivial source of distortion in the observed contact statistics. The protocol registers a contact only when two loci are successfully cross-linked, digested, and ligated into a chimeric fragment (Fig. 1D,E). Inefficient cross-linking or incomplete digestion effectively enlarge the capture radius *r*_*c*_, thereby reducing the apparent resolution of contacts at short genomic separations. We show that random digestion by restriction enzymes naturally leads to a Gaussian detection kernel 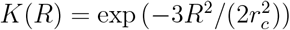, which reflects the probability that two loci separated by vector *R* are captured (see SI). The mean fragment length underlying this kernel is related to the capture radius by

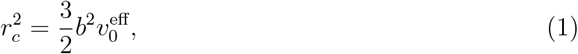

where 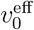 incorporates both restriction-fragment statistics and cross-linking efficiency. This effective parameterization allows Hi-C detection effects to be folded seamlessly into the polymer model.

Combining ideal-chain statistics with this Gaussian detection kernel yields the baseline Hi-C signal *P*_0_(*s*) in the absence of loops. For large separations 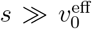, protocol effects become negligible and the familiar ideal-chain scaling *P*_0_(*s*) ∼ *s*^−3*/*2^ is recovered [16, 17]. In the opposote limit, 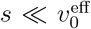, the entire segment of length *s* lies within the capture radius, producing a plateau in *P* (*s*) and a vanishing slope. The exact crossover is

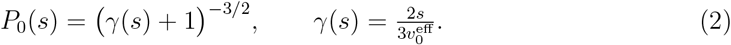

illustrating that protocol effects dominate short-range signal at *s* ∼ 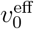, while leaving the log-derivative at larger scales essentially unchanged.

### Chromosomes with loops

We now turn to the wild-type situation, in which cohesin actively extrudes loops. To incorporate this, we use the quenched loop–disorder framework [6], modeling chromatin as a polymer decorated with a disordered array of static loops and gaps. Each loop is represented by a cohesin clamp bridging two non-adjacent loci, while intervening gaps correspond to unlooped polymer segments. Loop and gap lengths are drawn independently from exponential distributions with mean values *λ* and *g*, respectively, generating a stationary loop–gap pattern characteristic of steady-state loop extrusion (Fig. 1B).

At short genomic separations (*s* ≪ *λ, g*), the influence of loops can be computed perturbatively as a correction to the loop-free baseline in Eq. (2). To leading order, this yields

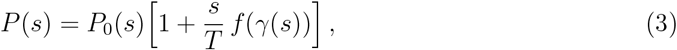

where *T* = *λ*+*g* is the loop–gap period (the inverse of the loop density), *f* (*γ*) = 3*γ/*[2(*γ* +1)], and *γ*(*s*) is the protocol-dependent scaling parameter defined in Eq. (2).

Importantly, the perturbative correction in Eq. (3) depends only on the loop period *T*, not on the loop and gap lengths separately. To understand this in simple terms, let us consider a genomic interval of length *s* placed randomly along the alternating loop–gap pattern (Fig. 1B), which yields three possible configurations:

a. **Gap–gap:** Both loci lie within the same gap. Their spatial statistics match those of a free polymer segment, contributing ((*g* − *s*)*/T*) *P*_0_(*s*), where (*g* − *s*)*/T* is the configuration probability.
b. **Loop–gap:** One locus lies inside a loop and the other in the adjacent gap. The probability of straddling a loop–gap boundary scales as *s/T*, since it is proportional to the length of the interval. Spatial statistics remain essentially free to leading order, yielding a contribution proportional to (*s/T*)*P*_0_(*s*).
c. **Loop–loop:** Both loci lie within the same loop. This occurs with probability (*λ*−*s*)*/T*, giving a zeroth-order contribution (*λ/T*)*P*_0_(*s*). At linear order, the loop constraint modifies the contact probability by a contribution proportional to (*s/λ*)*P*_0_(*s*); multiplying by the configuration weight cancels the factor of *λ*, again producing a term proportional to (*s/T*)*P*_0_(*s*).

Thus, the zeroth-order contributions from configurations (a) and (c) sum to the baseline *P*_0_(*s*), while the linear-order corrections from all three cases (a)-(c) yield the universal form ∼ (*s/T*)*P*_0_(*s*). This reduction explains why the short-scale folding of a looped polymer depends solely on the loop density. Taking the logarithmic derivative of Eq. (3) yields the central result of our theory:

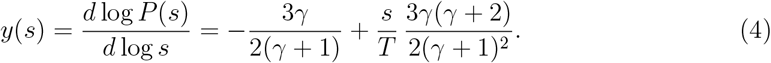

The first term in Eq. (4) reflects the effect of the finite capture radius alone (Eq. 2), whereas the second term represents the correction introduced by loop formation. The interplay between these two contributions gives rise to the characteristic local minimum—the “dip”—in the log-derivative curves (Fig. 2A; red theory curve in Fig. 2B). Neither loops nor finite detection alone can reproduce this feature, as demonstrated by theory (orange curve in Fig. 2B) and by cohesin-depleted Hi-C data, which display a monotonic decay of the log-derivative at tens of kilobases with no discernible dip (Fig. S7B). Likewise, multiple Micro-C datasets [13, 18] (Fig. S12) with capture radii as small as nucleosome spacing exhibit no detectable minimum, consistent with model predictions for a looped polymer in the limit of vanishing *r*_*c*_ (blue curve, Fig. 2B) [6].

The dip lies between the two genomic length parameters controlling the log-derivative at short scales: loop–gap period *T* and the effective fragment length 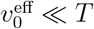 . Accordingly, the the dip position scales as the geometric mean of the two:

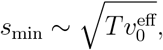

while its dimensionless depth *y*(*s*_min_) depends only on their ratio 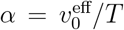. Increasing effective fragment length shifts the dip to larger scales and makes it shallower, behavior reproduced by perturbation theory and full quenched-loop calculations (Fig. S2). Experimentally, minima from diverse Hi-C datasets collapse along a diagonal band consistent with a conserved loop density across mammalian cell types (Fig. 2A).

Together, these results demonstrate that the short-scale dip in the log-derivative of *P* (*s*) emerges from the combined effects of cohesin-mediated looping and the finite detection radius inherent to Hi-C. This feature depends solely on loop density and experimental resolution, establishing a widely conserved fingerprint that recurs across mammalian datasets.

### Active-extrusion simulations validate equilibrium theory

Our theoretical framework treats loops as static, assuming that each cohesin-induced configuration thermally equilibrates. Although loop extrusion is intrinsically non-equilibrium, at scales less than the loop size this dynamical character becomes unimportant. Indeed, short segments thermally relax faster (quadratically in genomic length scale, *t* ∼ *s*^2^) than extrusion acts (linear, *t* ∼ *s*), yielding a crossover scale of *s*^*^∼0.5–1 Mb (see Methods). Accordingly, below *s < s*^*^, polymer segments have enough time to relax and remain equilibrated.

To support this scaling argument we additionally tested the theory predictions against molecular-dynamics simulations of chromatin undergoing *active* loop extrusion. Cohesins were modeled with realistic two-sided extrusion kinetics [5], parametrized by their mean spacing *d* (inverse linear density), processivity *l*, and extrusion rate *r* (see Methods for details).

### Phantom chain benchmark

We first examined loops on random-walk polymer chains without any persistence (Fig. 3A,B; Fig. S6). Strikingly, the log-derivatives from perturbation theory developed for static loops (Eq. (4)) quantitatively describe the curves for actively extruding system from simulations at short scales, provided the cohesin spacing *d* is identified with the loop period *T* . In this regime, the log-derivative depends only on loop density and fragment length 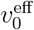, while the processivity *l* is irrelevant (Fig. 3B). The influence of *l* emerges only at larger scales, where a peak appears near the typical loop size *λ*(*l*) [6]. For different values of *d* (160 and 320 kb), perturbation theory and simulations match closely at short separations (Fig. S6), confirming that static-loop theory provides an accurate description of the active system at short genomic scales.

**Figure 3.**
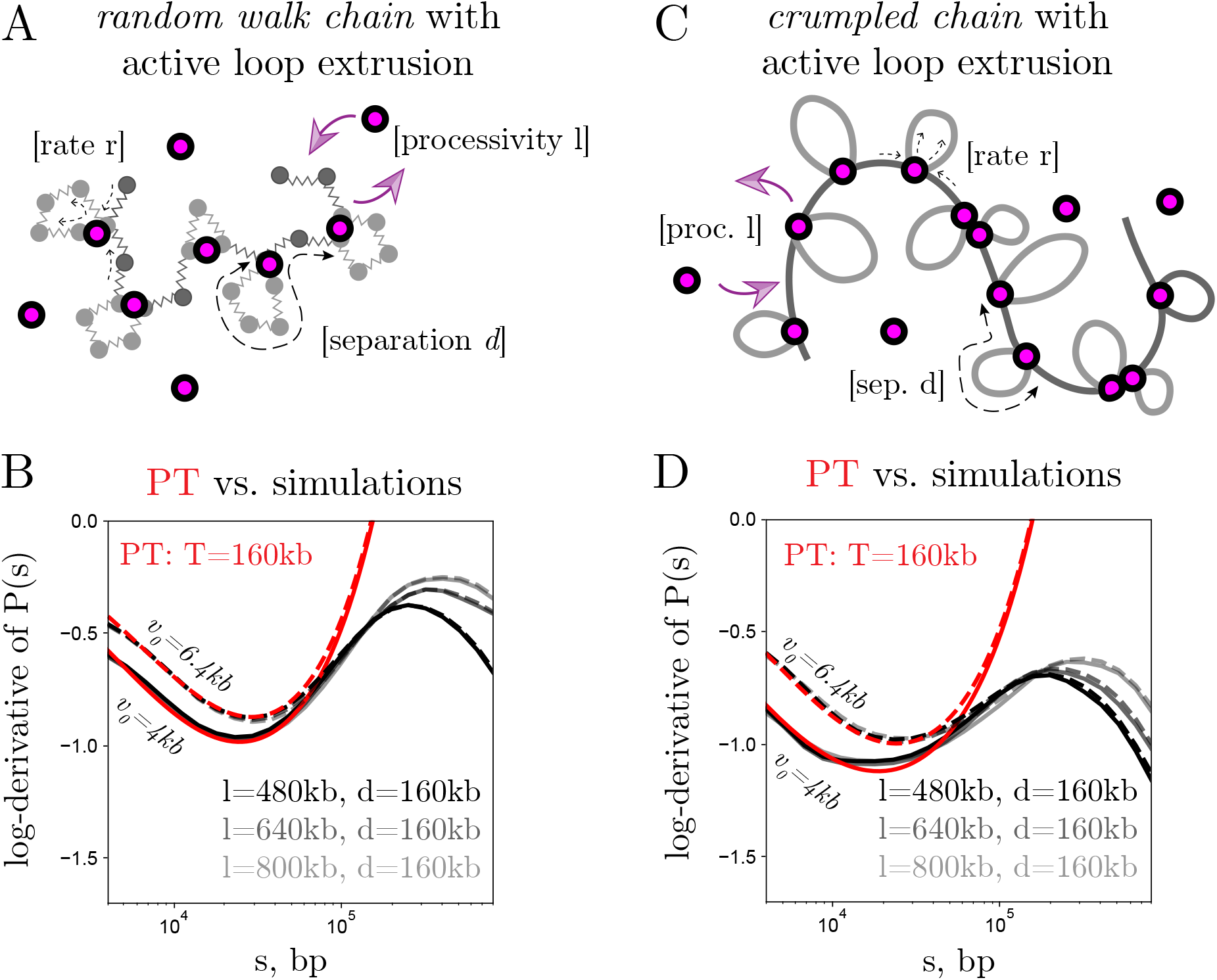
Validation of perturbation theory using molecular dynamics simulations of active loop extrusion. Simulations used cohesin spacing *d*, processivity *l*, extrusion rate *r*, and restriction fragment length *v*_0_. **(A)** Active loop extrusion on a phantom (random-walk) polymer. **(B)** Perturbation theory (PT) matches phantom-chain simulations when *T* = *d* and 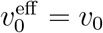; curves shown for *v*_0_ = 4 kb (solid) and 6.4 kb (dashed). **(C)** Active extrusion on a crumpled, semi-flexible polymer. **(D)** PT matches crumpled-chain simulations when *T* = *d* and 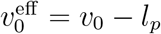 with *l*_*p*_ = 1.8 kb; *v*_0_ = 4 kb (solid) and 6.4 kb (dashed). One bead corresponds to 0.8 kb of chromatin.

### Crumpled chains and persistence correction

We next validated the theory in a more realistic chromatin context by simulating equilibrated melt of unknotted ring polymers [6, 12, 19], which reproduce the crumpled chromatin state observed in Hi-C and Micro-C experiments lacking cohesin [18]. Contact statistics from the baseline loop-free simulations were calibrated against Micro-C data without cohesin [18] to fix the conversion of 0.8kb/bead and persistence length, *l*_*p*_ = 1.8 kb (see Fig. S5). In this crumpled setting, perturbation theory again quantitatively matched the short-scale behaviour of log-derivatives from active extrusion simulations (Fig. 3D; Fig. S5), provided the effective fragment length 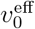is reduced by *l*_*p*_ (see SI). This correction accounts for the semi-flexibility of chromatin: fragments shorter than *l*_*p*_ behave as rigid, impermeable blobs and register essentially no contacts in the vicinity of their spatial extent. Crucially, the loop period *T* in this comparison remained unchanged.

Notably, the persistence correction explains the absence of a dip in Micro-C datasets (Fig. S12), where the restriction fragment length approaches the nucleosome spacing—well below the chromatin persistence length—yielding an effectively vanishing capture radius. This places Micro-C in the 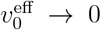 limit of our theory, where the dip is predicted to disappear.

Together, these simulations establish that equilibrium perturbation theory remains valid under non-equilibrium active extrusion with chromatin persistence absorbed in 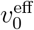. This result is non-trivial: even though cohesins are dynamically loading and extruding, short genomic segments relax faster than extrusion occurs, leaving a quasi-equilibrium signature in *P* (*s*).

### Experimental validation of the theoretical framework

To test our theoretical predictions, we analyzed two classes of experimental perturbations that selectively modulate either cohesin abundance or Hi-C resolution.

We first examined a recent Hi-C dataset in mouse embryonic stem cells (mESCs) from Shah *et al*. [20], in which the chromatin-bound cohesin fraction was systematically reduced using an auxin-inducible RAD21 degron system. The small molecule dTAG-13 was titrated at concentrations of 20 and 45 nM to induce partial degradation of RAD21 and thereby reduce cohesin occupancy on chromatin. Flow cytometry confirmed that approximately 50% of cohesin remained chromatin-bound at 20 nM and about 30% at 45 nM dTAG.

We asked whether these measured reductions in cohesin abundance could be independently reproduced by our theoretical inference. For each degron concentration, we computed the short-scale logarithmic derivative of the contact probability *P* (*s*) (Fig. 4A). Increasing dTAG concentration produced a systematic rightward and downward shift of the dip—corresponding to larger *s*_min_ and smaller *y*_min_—consistent with an increased loop period *T* and reduced loop density *T* ^−1^. Fitting the data using Eq. (4) revealed a decrease in loop density from 10 loops/Mb in wild type to 5.6 loops/Mb (56% remaining) at 20 nM and 4.5 loops/Mb (45% remaining) at 45 nM, closely matching the cohesin retention from flow cytometry (50% and 30%, respectively) [20]. Notably, the fitted effective fragment length, 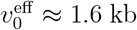, remained nearly constant across all concentrations, confirming that the observed changes in *P* (*s*) are explained solely by altered cohesin abundance.

**Figure 4.**
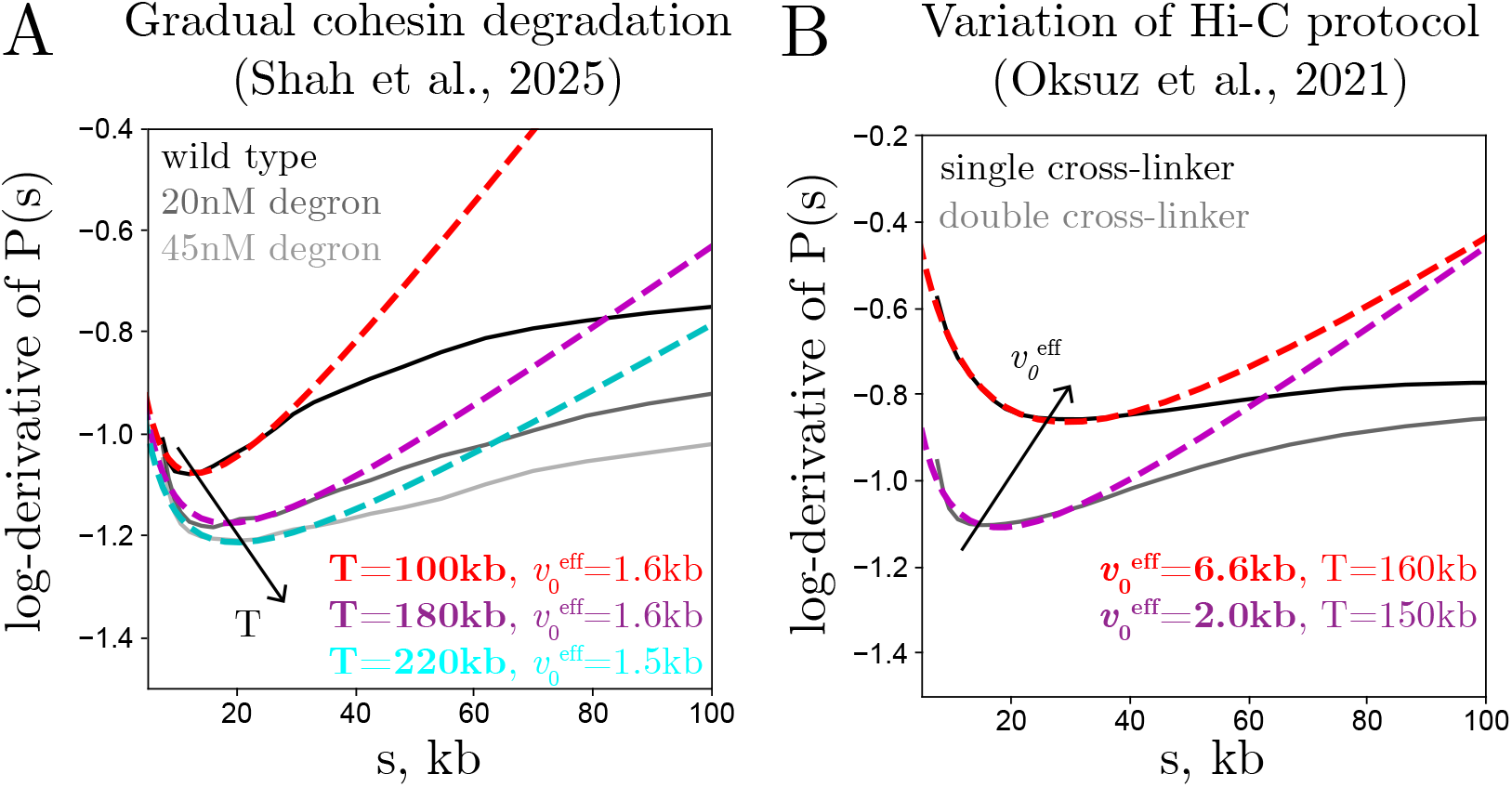
Experimental validation of the theoretical framework. **(A)** RAD21 degron titration (20, 45 nM) in mESCs reduces loop density from ∼10 to ∼4.5 loops/Mb, while 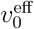remains constant. Curves show best-fit theory (Shah *et al*. [20]). **(B)** Enhanced cross-linking (FA+DSG) in HFF cells decreases 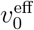 by ∼3-fold relative to standard in situ Hi-C, with loop density largely unchanged (∼6.5 loops/Mb) (Oksuz *et al*. [13]).

We further analyzed another perturbation targeting the cohesin loader NIPBL in the same system [20]. NIPBL promotes cohesin loading onto DNA and may also facilitate its extrusion rate. Fitting this profile against the theoretical model (Fig. S4A) yielded a loop period of *T* ≈ 160 kb, corresponding to a loop density of 6.3 loops/Mb (∼63% of wild-type levels). This reduction mirrors the effect of partial RAD21 degradation and agrees with independent flow cytometry measurements showing that ∼50% of cohesin remains DNA-bound after NIPBL degradation [20]. Together, these analyses demonstrate that our framework quantitatively captures changes in loop density arising from either direct cohesin degradation or impaired loading efficiency.

To assess an orthogonal perturbation affecting experimental resolution rather than cohesin levels on chromosomes, we analyzed Hi-C data from Oksuz *et al*. [13]. In this study, cross-linking and fragmentation conditions were systematically varied while maintaining identical chromatin organization. We compared standard in situ Hi-C (FA cross-linking, DpnII digestion) to the enhanced Hi-C 2.0 protocol employing dual cross-linkers (FA+DSG, DpnII digestion). As shown in Fig. 4B, the dual cross-linking condition caused a leftward and downward shift of the dip in HFF cells, reflecting a decrease in 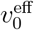 consistent with higher capture efficiency. Theoretical fits reproduced both curves accurately, revealing a nearly threefold reduction in 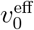 upon addition of DSG, while the loop period remained nearly unchanged. A similar ∼3-fold reduction in 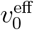 was observed in the hESC system for the equivalent addition of cross-linker (Fig. S4B), with loop density change less than by 20%. Together, these two complementary perturbations—one altering cohesin density and the other contact capture resolution—provide strong experimental validation of our theoretical framework and demonstrate its ability to quantitatively extract key physical parameters from Hi-C contact statistics.

### Quantifying loop density across datasets and inferring cohesin stoichiometry

Our polymer framework establishes a direct quantitative connection between a universal short-scale Hi-C signature and the density of cohesin-mediated loops. To systematically assess the parameters across systems, we compiled the positions and depths of the log-derivative minima (*s*_min_, *y*_min_) from 33 Hi-C datasets spanning both human (16) and mouse (17) experiments that differ in cell type, developmental state, and protocol (from Fig. 2A; STable 1). Each dataset was compared to theoretical trajectories for varying loop periods *T* (Fig. 5A). Remarkably, all data points fall along the theoretical family of curves corresponding to *T* = 100–250 kb, enabling robust parameter extraction for each dataset.

**Table 1.**
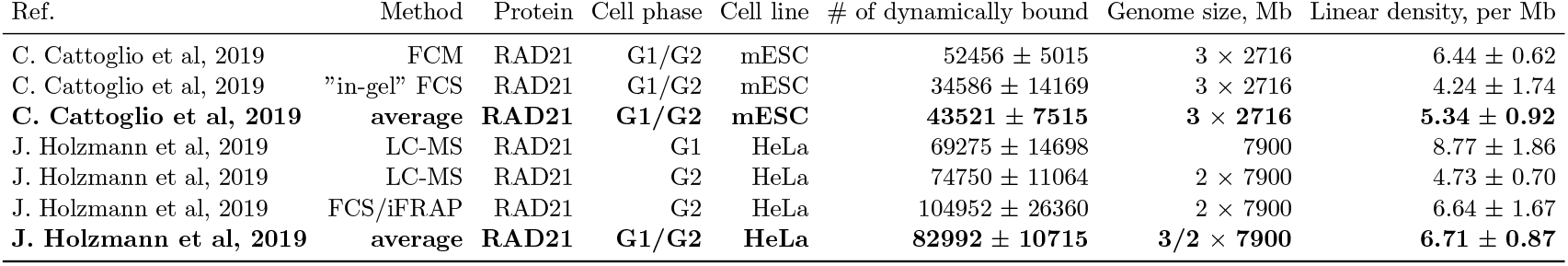
Cohesin abundance estimates across cell types and measurement methods, compiled from Cattoglio *et al*. [9] and Holzmann *et al*. [26]. Bold entries denote averages across methods within each study. Linear densities were computed from dynamically chromatinbound RAD21 counts. STED-based values from [11] are excluded, as they likely reflect concentration-dependent clustering rather than true copy numbers.

**Figure 5.**
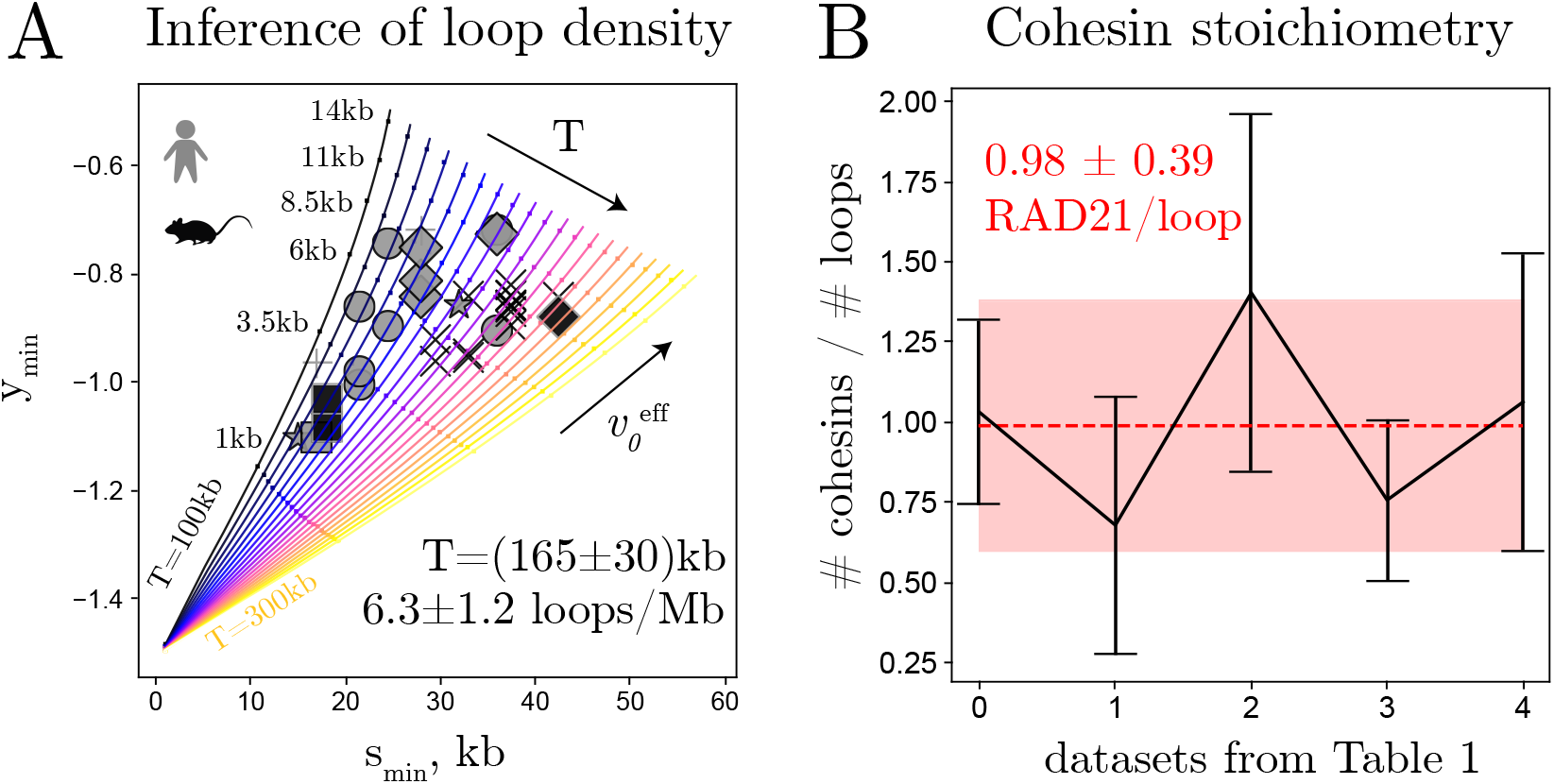
Quantitative inference of cohesin loop properties from human and mouse Hi-C datasets (same as Fig. 2A). **(A)** Positions (*s*_min_) and depths (*y*_min_) of log-derivative minima overlaid on theoretical trajectories for loop periods *T* = 100–300 kb; 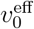 increases along each curve. Across datasets, the mean inferred period is *T* = 165 30 kb (6.25 *±*1.15 loops/Mb). **(B)** RAD21 copy numbers (Table 1) compared with inferred loop counts. After genomic normalization, cohesin abundance closely matches loop number, consistent with approximately one RAD21 complex per loop.

We find that while the inferred effective fragment lengths 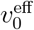 vary substantially across experiments (from ∼ 1 to ∼ 10 kb), reflecting differences in the experimental protocol (STable 1), the loop periods are narrowly distributed around

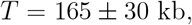

corresponding to a conserved cohesin-mediated loop density of 6.25*∼*1.15 loops per megabase. The modest (∼20%) uncertainty reflects striking consistency across cell types, organisms, and experimental conditions. Importantly, the inferred loop density is reproducible across biological and technical replicates (Fig. S9). This result suggests that loop density is a fundamental and conserved feature of mammalian interphase chromatin.

Notably, our inference complements previous modeling efforts that estimated loop parameters by fitting large-scale Hi-C features such as TAD structures or global *P* (*s*) slopes, yielding comparable values (4-8 loops/Mb) [5, 6, 21–25]. By focusing on short genomic separations, here we avoid the degeneracies inherent to large-scale fits and extract loop density in a parameter-complete, robust manner.

### Stoichiometry of extruding cohesion

The fact that the dip shifts progressively with partial RAD21 degradation and disappears upon cohesin depletion indicates that the dip-inferred loop density reflects the abundance of cohesin-mediated loops. This enables a direct comparison between the loop densities inferred from Hi-C and the densities of chromatin-bound cohesin complexes measured independently by quantitative fluorescence spectroscopy and mass spectrometry (Table 1). Across systems, these two quantities agree closely, yielding an average of 0.98 *±* 0.39 cohesins per inferred loop (Fig. 5B). Reported RAD21 densities of 5–7 cohesins per megabase in human and mouse cells [9, 26] thus fall within the range expected from the ∼6 *±* 1 loops/Mb inferred from Hi-C. This correspondence suggests that, on average, a single cohesin complex is sufficient to extrude a loop under interphase conditions, arguing against widespread dimeric extrusion.

As an important caveat, a recent super-resolution imaging study reported substantially higher apparent cohesin copy numbers on HeLa cells (11–12 per Mb) [11], which—if taken at face value—would imply dimeric extrusion. However, as argued by the authors of the same study, these elevated figures, roughly twice those obtained previously in the same system [26], arise from concentration-dependent clustering of the abundant STAG2 isoform in the course of interphase progression, which was overlooked previously. Such clustering increases local cohesin encounters and can contribute to the appearance of paired complexes [10, 11] without true stoichiometric dimerization. Consequently, these inflated counts likely reflect aggregation of unbound cohesin with loop-extruding molecules rather than genuine extrusion dimers. We therefore caution against using these values directly in stoichiometric estimates. Overall, the close quantitative correspondence between inferred loop numbers and absolute cohesin counts across datasets and organisms provides genome-wide support for predominantly monomeric cohesin extrusion during interphase. Notably, our analysis constrains the *average* extrusion stoichiometry to be predominantly monomeric, but cannot exclude rare or transient dimeric interactions that may occur locally or under specific conditions.

### Dynamics of chromatin loop reorganization during mitotic exit

We next applied the framework to a dynamic biological transition—mitotic exit—during which condensins dissociate and cohesins reload to progressively establish interphase chromatin architecture [11, 27]. To this end, we analyzed synchronized Hi-C time courses from Abramo *et al*. (HeLa) [28] and Zhang *et al*. (mouse erythroblasts GE1) [29] (Fig. 6; Figs. S10–S11).

**Figure 6.**
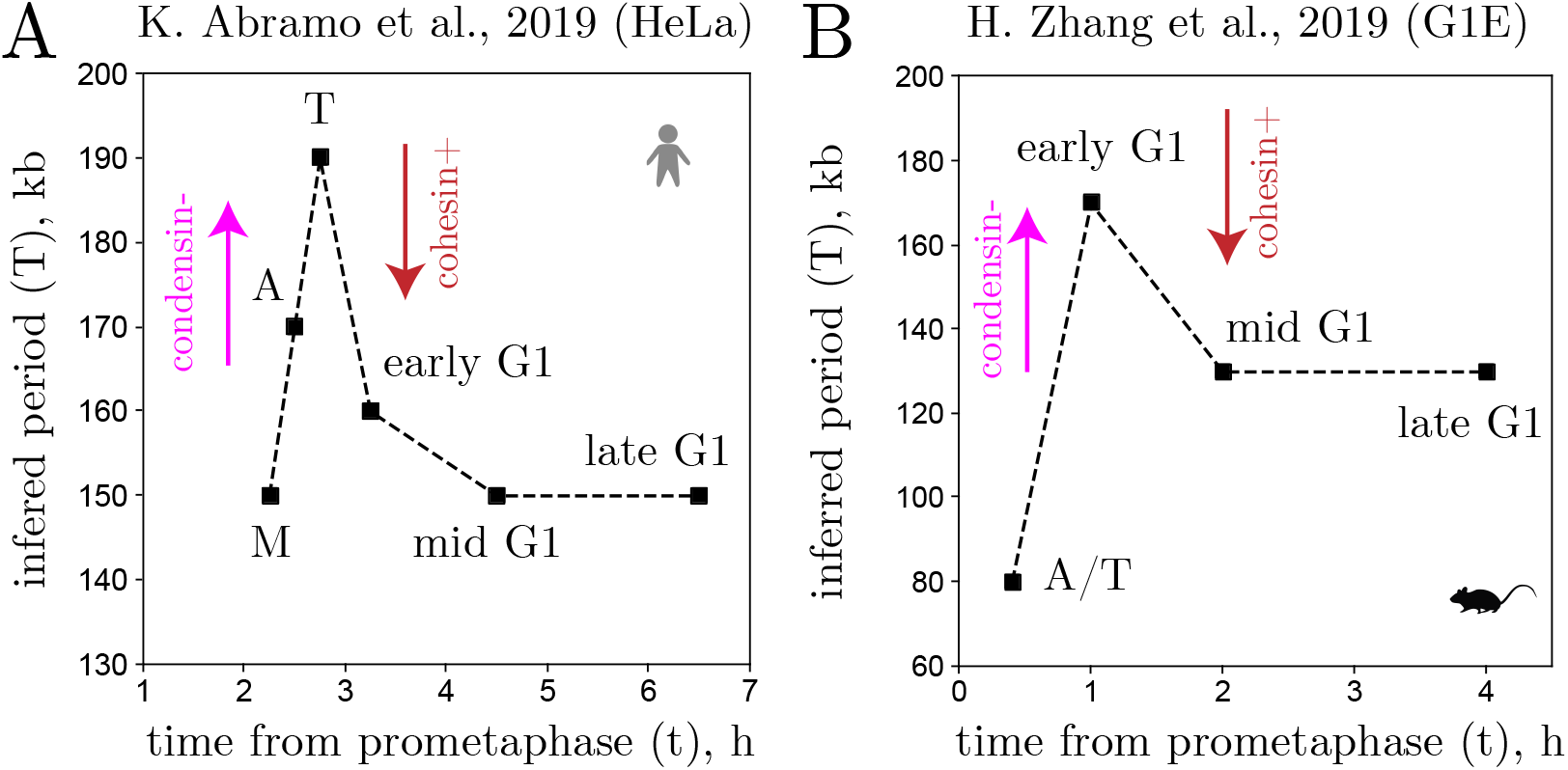
Chromatin loop reorganization during mitotic exit from Hi-C time-course data. Shown are inferred loop periodicities *T* (*t*) from Abramo *et al*. [28] (A) and Zhang *et al*. [29] (B). Both datasets exhibit a transient increase in *T* as condensins unload during mitotic exit, followed by a decrease as cohesin reloads to establish interphase loop architecture. Time *t* denotes hours after release from prometaphase arrest.

Tracking the positions of log-derivative minima revealed pronounced temporal dynamics in loop periodicity in both datasets. In the HeLa system (Abramo *et al*.), loop spacing peaked at telophase, indicating a transient reduction in loop density during the condensin–cohesin handoff, consistent with the existence of an intermediate folding state at this stage [28]. In contrast, in the Zhang dataset, the largest loop period occurred later, at early G1, with the highest loop density detected in anaphase. This pattern is consistent with the slower binding of cohesin in that study: RAD21 ChIP–seq profiles indicated minimal levels of cohesin occupancy until early G1 [29]. These complementary trends align with recent super-resolution imaging of HeLa cells [11], which revealed rapid loading of STAG1-isoform of cohesin immediately after telophase, followed by gradual STAG2-isoform recruitment. Together, these results suggest that the precise kinetics of cohesin accumulation differs modestly between human and mouse systems, reflecting species- or cell-type–specificities. The observed trajectories in both systems capture the sequential disengagement of condensin and loading of cohesin, providing an independent validation of the framework’s applicability to dynamic chromatin reorganization in the cell cycle.

## Discussion

We present a quantitative polymer-physics framework that addresses a central mechanistic question in genome biology: the linear density of cohesin-mediated loops along interphase chromatin. By combining a quenched loop-disorder approach with a finite capture radius of Hi-C, we construct a perturbation theory for the short-range contact probability *P* (*s*) and its log-derivative. The theory predicts a characteristic dip in the logarithmic derivative, arising jointly from the loops and the finite Hi-C resolution. Neither effect alone can reproduce this feature. The ubiquity of this dip across mammalian Hi-C datasets provides additional evidence for genome organization through cohesin-extruded loops in living cells.

From a polymer-physics perspective, short-range statistics of a looped chain reduce to a single structural parameter—the loop period—making the position of the log-derivative minimum depend only on the linear density of loops and the Hi-C capture radius. This prediction is confirmed by molecular dynamics simulations of state-of-the-art active extrusion, where the separation between motors, rather than processivity, governs short-scale statistics (Fig. 3). Experimental perturbations provide independent validation: partial cohesin degradation or impaired loading produces a gradual, concentration-dependent decrease in inferred loop density (Fig. 4A, S4A), whereas protocol enhancements in Hi-C 2.0 leave loop density unchanged but reproducibly reduce the capture radius by a factor of 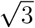 (Fig. 4B, S4B), providing a quantitative handle on resolution effects. These results establish that short-range Hi-C statistics offer an experimentally accessible fingerprint of cohesin loop density and can be also used to monitor dynamic chromatin reorganization (Fig. 6).

Application of the inference framework to mammalian interphase datasets reveals a conserved loop density of 6.3 *±* 1.2 loops per megabase, corresponding to loop periods *T* ≈ 170 kb (Fig. 5A). Assuming a mean loop size of *λ* ≈ 100–120 kb [6, 11, 12], implies typical gap lengths of *g* ≈ 50 − 70 kb and that roughly *λ/T* ≈ 60–70% of the genome is contained within cohesin-mediated loops. Remarkably, a recent live-imaging study at the *Fbn2* domain reported that 57–61% of chromatin resides within loops [30], in striking agreement with our genome-wide inference.

Comparing inferred loop densities with absolute cohesin abundances measured by quantitative imaging and mass spectrometry (Table 1) reveals near-perfect agreement—approximately six cohesin complexes per megabase, or roughly one cohesin per loop— supporting a predominantly monomeric mode of loop extrusion in vivo. At first glance, this conclusion appears to contrast with recent super-resolution microscopy, which reports that extruding (non-cohesive) cohesin complexes often appear as closely paired particles, whereas cohesive complexes maintaining sister chromatids in G2 are clearly monomeric [10, 11]. However, as noted by the authors of [11], these paired signals likely arise from concentration-dependent clustering of the abundant STAG2 isoform, which can mimic the appearance of extrusion dimers without reflecting true stoichiometric pairing. Crucially, the less abundant and less aggregation-prone STAG1 isoform—also capable of loop extrusion—was consistently observed as a spatially isolated monomer, further reinforcing the view that the vast majority of extrusion events are mediated by single cohesin complexes.

Across datasets, the second inferred parameter—the effective fragment length 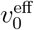 characterizing the capture radius —ranged from 1–10 kb (mean 6.4 kb; STable 1), consistent with restriction fragment sizes in standard Hi-C protocols [13]. However, 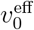 should be viewed as a phenomenological parameter reflecting both physical factors (chromatin stiffness) and biochemical aspects of the Hi-C workflow, including cross-linking chemistry [13], multi-ligation events [31], and random ligation of diffusing fragments [14]. Using the Gaussian relation 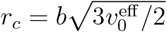 with *b* ≈ 0.05*µm* yields a typical capture radius of *r*_*c*_ ≈ 0.15*µm*. Interestingly, this number is comparable to the chromatin mesh size ξ ∼ (*V/L*)^1*/*2^, where *V* ∼ (10 *µ*m)^3^ is nuclear volume and *L* ∼ 1 m is the total chromatin length [19]. This correspondence supports the idea that topological constraints at the chromatin mesh limit diffusion of crosslinked restriction fragments and constrain the scale of the contact capture.

In summary, a two-parameter physical model of looped polymers captures a universal feature of short-range Hi-C contact statistics and links it quantitatively to cohesin abundance. By bridging polymer physics with molecular stoichiometry, this framework provides a robust, interpretable means to infer loop density and motor organization directly from population Hi-C maps. Beyond investigating the cohesin monomer–dimer controversy, it establishes short-range Hi-C contact statistics as a sensitive quantitative probe of the molecular mechanisms shaping genome architecture.

## Data and Code Availability

All preprocessed datasets analyzed in this study, together with Jupyter notebooks reproducing the main figures and a standalone Python script for parameter inference from experimental Hi-C scalings, are publicly available at the project’s GitHub repository: https://github.com/kipolovnikov/loop-density-hic.

## Methods

### Scaling estimate of the crossover scale

A sufficiently short genomic segment (*s < s*^*^) achieves thermal equilibration faster than cohesin can perturb its spatial organization by active loop extrusion. The crossover length *s*^*^ follows from comparing the Rouse relaxation time *τ*_*R*_(*s*) ∼ *τ*_0_(*s/l*_*K*_)^2^ (with *τ*_0_ the Kuhn-scale *l*_*K*_ diffusion time) to the extrusion time *τ*_ex_(*s*) = *s/r*, where *r* is the extrusion rate. Using chromatin estimates *τ*_0_ ∼ 10^−2^ s [6, 32], *l*_*K*_ ∼ 4 kb [15, 33], and *r* ∼ 1–2 kb/s [7, 8], we find the crossover scale *s*^*^ ≈ 0.5–1 Mb. Thus, at the sub-loop scales (*s < λ, g* ≈ 50 − 100 kb) central to our study, the equilibrium assumption is well justified.

### Molecular dynamics simulations of active loop extrusion

All simulations were performed with the `polychrom` framework (https://github.com/mirnylab/polychrom), which implements Langevin dynamics for bead–spring polymers. Chromatin fibers were modeled as chains of *N* = 40,000 beads connected by harmonic springs. One bead was mapped to 800 bp by comparison of the crumpled chain simulation with Micro-C data (see SI); this conversion of 0.8kb/bead was used for both phantom and crumpled chains.

### 1D extrusion model

Two-sided loop extrusion was implemented by coupling 3D polymer dynamics to a 1D stochastic model of cohesin kinetics [5, 34]. Each cohesin was represented as an additional harmonic bond between two beads. At initialization, *N*_0_ = *N/d* cohesins were placed randomly along the chain, defining the mean spacing *d*. During simulations, each bound cohesin unbound with rate *k*_off_ = 2*r/l*, where *l* is the processivity length and *r* the extrusion speed, and the same number of cohesins were reloaded at random unoccupied sites to maintain *N*_0_ constant. Upon binding, cohesins extruded symmetrically at speed *r* ≈ 1 kb/s until unbinding, collision with another cohesin, or reaching the chain ends. No extrusion barriers (e.g. CTCF, RNA polymerase) were included, to isolate the fundamental effect of cohesin motors. Simulation time was mapped to real time by calibrating the Rouse diffusion coefficient of individual beads, yielding *D*_*R*_ ≈ 10^−2^ *µ*m^2^ s^−1*/*2^, consistent with experimental measurements [30, 35].

## Acknowledgements

We gratefully acknowledge Job Dekker and Leonid Mirny for valuable feedback on the manuscript. This work was supported by the Russian Science Foundation (Grant No. 25-13-00277). K.P. also acknowledges support from the Alexander von Humboldt Foundation.

