## Supplementary Materials for "A universal polymer signature in Hi-C resolves cohesin loop density and supports monomeric extrusion"

#### Contents

**1 Introduction** **2**

**2 Quenched disorder of random loops** **3**

**3 Spatial statistical properties** **4**

**4 Statistical weights** **5**

**5 Capture radius models** **6**

**6 Perturbation theory** **10**

**7 MD simulations** **15**

**8 Hi-C data analysis** **22**

### 1 Introduction

This appendix (i) complements the minimal loops-plus-cutoff-radius model of interphase chromatin developed in the main text, (ii) provides the full mathematical derivations needed to analytically describe the short-scale behavior of the contact probability $P(s)$ in the presence of quenched loop disorder and a finite Hi-C detection radius, (iii) provides the detailed analysis of molecular dynamics simulations and (iv) provides the description and analysis of various Hi-C datasets.

Section 2 introduces the model of quenched loop disorder in which the polymer is represented as an alternating sequence of non-overlapping loops of length $\lambda$ and gaps of length $g$. We formalize how this alternation induces a small set of mutually exclusive geometric configurations for any pair of loci and set up the integral representation of $P(s)$ as a sum over these configurations.

Section 3 recalls the spatial statistics of a Gaussian polymer with fractal dimension $d_f$ and spells out the end-to-end variances corresponding to each configuration. These expressions determine the spatial kernels that enter the contact integrals. Section 4 then provides the probability densities (statistical weights) with which the configurations occur along the contour, derived from the two-state loop/gap alternation. The result is a closed set of weights and variances that, together, control the disorder-averaged distribution of 3D separations at fixed contour distance $s$.

Building upon the framework of Sections 2–4, Section 5 incorporates the capture radius into the modeling. Subsections 5.1–5.3 present several complementary formulations for contact detection, namely a Gaussian capture kernel and two fragment-aware models. Subsection 5.4 establishes the correspondence between these formulations, outlines the parameter regimes in which they are interchangeable, and isolates the effective capture radius that can be carried through the remaining analysis.

Section 6 exploits the small-scale limit $s \ll \lambda, g$ to construct a perturbative expression for $P(s)$ and for its logarithmic derivative $y(s) = \mathrm{d}\log P(s)/\mathrm{d}\log s$. It provides explicit derivation of formulas used in the main text for parameter inference and interpreting short-range features of Hi-C contact profiles.

Finally, Section 7 provides the detailed discussion of the results of molecular dynamics simulations and Section 8 contains the details of Hi-C datasets used in the paper.

#### 2 Quenched disorder of random loops

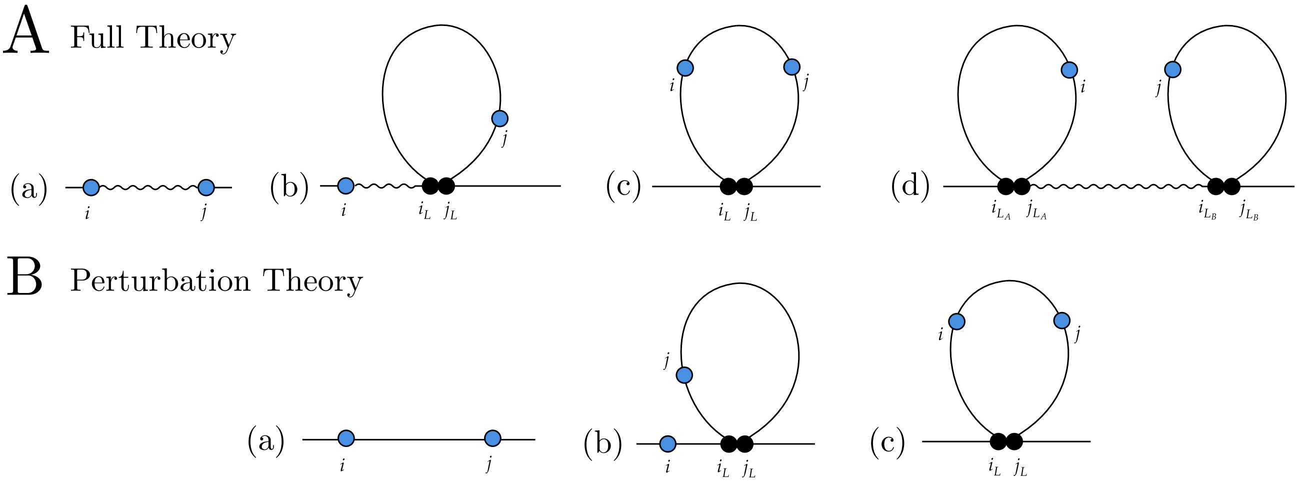

Figure S1: (A) Diagrams that fully describe possible linear positions of contact points $i$ and $j$ in the chain with alternating loops and gaps. Solid lines represent a specific loop or gap segment, whereas wavy lines represent a segment with arbitrary number of loops and gaps in-between. (B) A subset of those diagrams that give a linear contribution to contact probability in the limit $s/\lambda \ll 1$ and $s/g \ll 1$, see Eq. (6.7).

We start by introducing the model of *quenched loop disorder*. Within this framework, the polymer chain relaxes fast enough so that each distinct topological conformation can be considered equilibrated - in other words, we average with respect to thermal noise before performing averaging with respect to configuration disorder (see the main text for justification). Each such configuration (which we term *diagrams*) consists of two *alternating* elements - loops, whose contour lengths are distributed exponentially with mean $\lambda$, and gaps between them, whose contour lengths are distributed exponentially with mean $g$, thus constituting the linear structure of any chain conformation in our model. We stress that there are no overlapping or nested loops in our model - each loop is followed by a gap, and each gap is followed by a loop.

Afterwards, we aim to calculate the expression for spatial contact probability $P(s)$ between two points of interest with contour coordinates $i$ and $j$, as a function of their linear separation $s = |j - i|$. We do so by considering contributions to total probability from several diagrams, representing different positions of the loops with respect to contact points, see Fig. S1A. Formally speaking, before making any assumptions about equilibrium spatial statistics of the chain, probability of an arbitrary *contact event* in this model can be calculated using the law of total probability and consequent diagrammatic decomposition as follows

$$\begin{aligned} \mathbb{P}(\text{contact}) &= \int d\{A\}\ \mathbb{P}(\text{contact} \mid \{A\}) \cdot \omega(\{A\}) = \\ &= \sum_{i=(a,b,c,d)} \int_{B_i} d\{A\}\ \mathbb{P}^{(i)}(\text{contact} \mid \{A\}) \cdot \omega^{(i)}(\{A\}). \end{aligned} \tag{2.1}$$

In the first equality, we introduced two notions: a set of stochastic parameters $\{A\}$ that define the configuration of the loops with respect to contact points (for example, $\{A\}$ can be $\{i_L, j_L\}$, contour coordinates of the ends of the loop, or $\{i_{L_A}, j_{L_A}, i_{L_B}, j_{L_B}\}$ if there are two loops, or in practice some linear combination of those variables), and $\omega\{A\}$ is the *statistical weight* - the probability density of encountering such set of parameters in the alternating loop-and-gap ensemble. This relation arises due to the aforementioned law of total probability $\int d\{A\}\ \omega(\{A\}) = 1$ - that is, some configuration of stochastic coordinates occurs, and does so uniquely. In the second equality of Eq. (2.1) we separated the set $\{A\}$ into several distinct non-overlapping territories, $(a), (b), (c)$ and $(d)$, which correspond to diagrams from Fig. S1, by introducing integration volumes $B_i$ - the set of values that these parameters must take in relation to contour distance $s$ to constitute the diagram under the corresponding index $i$. Building upon this framework, we provide specific forms of $\mathbb{P}^{(i)}(\text{contact} \mid \{A\})$, $\omega^{(i)}(\{A\})$ and $B_i$ suitable for calculations in the following sections.

### 3 Spatial statistical properties

We model chromatin as a fractal chain with effective Kuhn length $b$ and fractal dimension $d_f$, whose end-to-end probability distribution is a zero-mean Gaussian characterized by variance $\sigma$

$$P(R \mid \sigma) = \frac{1}{(2\pi\sigma^2/3)^{3/2}} \exp\left(-\frac{3R^2}{2\sigma^2}\right), \tag{3.1}$$

which in the free case is equal to $\sigma^2 = \sigma_f^2(s) = b^2 s^{2/d_f}$, and $b \approx \sigma_f(l_K) l_K^{-1/2} \approx 0.05\mu m$ is the effective Kuhn length in spatial units, obtained by scaling the spatial extent of a Kuhn segment ($\sigma_f(l_K) \approx 0.1\mu m$) by its contour length $l_K \approx 4$kb. In our model, functional form of this distribution remains the same for every diagram, but variance $\sigma^2$ behaves differently from the unconstrained case. In order to obtain it for cases with loops, we have to use the assumption of non-topological attachment of loop extruders to the chain - that is, they grip rather than thread the fibre. More practically, this means that the chain consists of one independent backbone, and many independent loops which decrease its effective length without altering its equilibrium statistics. We will briefly recount how to calculate variances for all of these cases.

For diagram (a), variance is simply the same as the variance of a free fractal chain of length $s$, whose contour distance is effectively decreased by a factor $x$, which corresponds to the fraction of loops extruded inbetween end-points:

$$\sigma^2_{(a)}(s, x) = \sigma_f^2((1-x)s) = b^2((1-x)\cdot s)^{2/d_f} \tag{3.2}$$

For diagrams (b) through (d), we have to use the expression for variance of Brownian bridge of length $L$, which can be obtained using expressions for correlator of fractal Brownian motion and decomposition of variance of Gaussian vectors:

$$\sigma_l^2(s, L) = \sigma_f^2(s) \cdot \left(1 - \frac{(\sigma_f^2(L) + \sigma_f^2(s) - \sigma_f^2(L-s))^2}{4 \cdot \sigma_f^2(s) \cdot \sigma_f^2(L)}\right), \tag{3.3}$$

where $L$ is the length of the loop where contact points reside. Thus, for diagram (b) variance can be obtained as a sum of variances of gap and loop segments:

$$\sigma^2_{(b)}(s, l_1, l_2, x) = \sigma^2_f((1-x)\cdot(s-l_2)) + \sigma^2_l(l_2, l_1+l_2), \tag{3.4}$$

where $l_1 = j_L - j$, $l_2 = j - i_L$, and $x$ is the fraction of loops between $i$ and $i_L$. For diagram (c) we can simply use the expression for Brownian bridge, since both contact points reside on the loop:

$$\sigma^2_{(c)}(s, l_1, l_2) = \sigma^2_l(s, l_1+l_2), \tag{3.5}$$

where $l_1 = i - i_L$ and $l_2 = j_L - i$. Finally, for diagram (d) we have to sum three consecutive independent segments:

$$\sigma^2_{(d)}(s, l_1, l_2, h, L, x) = \sigma^2_l(l_2, l_1+l_2) + \sigma^2_f((1-x)\cdot h) + \sigma^2_l(s-h-l_2, L), \tag{3.6}$$

where $l_1 = i - i_{L_A}$, $l_2 = j_{L_A} - i$, $h = i_{L_B} - j_{L_A}$, $L = j_{L_B} - i_{L_A}$, and $x$ is the fraction of loops between $i_{L_B}$ and $j_{L_A}$. These four expressions fully define spatial part of the statistical properties of the chain, and will be used for the calculation of total contact probability.

### 4 Statistical weights

Turning our attention to statistical weights $\omega(\{A\})$ introduced in Section 2, we note that the system of exponentially distributed alternating loops and gaps can be mapped onto a two-state continuous Markov process with transition rates $\alpha_g = 1/g$ and $\alpha_l = 1/\lambda$. Using this analogy, we can extract two useful relations. The first one is the probability that the point at contour distance $s$ from the current one, which is known to be on a gap, lies on a gap as well:

$$g(s) = \frac{\alpha_l}{\alpha_g+\alpha_l} + \frac{\alpha_g}{\alpha_g+\alpha_l}\exp\left[-\left(\alpha_g+\alpha_l\right)s\right]. \tag{4.1}$$

The second is for the probability of encountering a segment, which, given that it starts and ends on a gap and has length $s$, also has a fraction of its contour $x$ occupied by loops:

$$\mathcal{F}(x \mid s) = \frac{\alpha_g+\alpha_l}{\alpha_l+\alpha_g e^{-(\alpha_l+\alpha_g)s}}\left\{e^{-\alpha_g s}\delta(x) + \left(\frac{\alpha_g\alpha_l(1-x)s^2}{x}\right)^{1/2} I_1\left[2\sqrt{\alpha_g\alpha_l x(1-x)s^2}\right] e^{-\alpha_g(1-x)s-\alpha_l x s}\right\}, \tag{4.2}$$

where $I_1$ is the modified Bessel function, $\delta(x)$ is Dirac delta function, and $x \in [0,1]$. We also introduce two auxilary functions $\pi_g = \alpha_l/(\alpha_g+\alpha_l)$ and $\pi_l = \alpha_g/(\alpha_g+\alpha_l)$, which are the probabilities that a randomly selected point lies on a gap or on a loop correspondingly. After obtaining these results, statistical weights of all the diagrams can be computed in a straightforward manner. For diagram (a) we get

$$\omega^{(a)}(x \mid s) = \pi_g \cdot g(s) \cdot \mathcal{F}(x \mid s), \tag{4.3}$$

which means that first we select a random point $i$ on a gap with probability $\pi_g$, then we multiply it by probability $g(s)$ of having its contact point $j$ lie on a gap as well, and then multiply it by the overall probability of encountering $x$ fraction of loops in the formed segment. The only restriction here is that $x \in [0,1]$ For diagram (b) we get

$$\omega^{(b)}(l_1, l_2, x \mid s) = 2 \cdot \pi_g \cdot g(s - l_2) \cdot \mathcal{F}(x \mid s - l_2) \cdot \alpha_g \cdot \rho(l_1 + l_2), \tag{4.4}$$

where the factor 2 accounts for the symmetrical diagram, $\pi_g$ is the probability to find $i$ on a gap, $g(s-l_2)$ is the probability to have $i_L$ on a gap, $\mathcal{F}(x \mid s-l_2)$ if the probability to encounter $x$ fraction of loops from $i$ to $i_L$, $\alpha_g$ is the probability to transition into a loop state, and the last factor $\rho(l_1+l_2)$ describes the probability of encountering a loop segment of given length, where $\rho(L) = \alpha_l \cdot \exp(-\alpha_l \cdot L)$. Restrictions for values of these parameters are $x \in [0,1]$, $l_1 \in [0,\infty)$ and $l_2 \in [0,s]$. For diagram (c) we get

$$\omega^{(c)}(l_1, l_2 \mid s) = \pi_l \cdot \rho(l_1) \cdot \rho(l_2), \tag{4.5}$$

which corresponds to the probability of finding a point on a loop which has partial lengths of $l_1 = i - i_L$ and $l_2 = j_L - i$. Their possible values are $l_1 \in [0,\infty)$ and $l_2 \in [s,\infty)$. Finally, for diagram (d) we get

$$\omega^{(d)}(l_1, l_2, h, L, x \mid s) = \pi_l \cdot \rho(l_1) \cdot \rho(l_2) \cdot g(h) \cdot \mathcal{F}(x \mid h) \cdot \alpha_g \cdot \rho(L), \tag{4.6}$$

which corresponds to the probability of finding point $i$ on a loop which has partial lengths $l_1$ and $l_2$, encounter a segment of length $h$ with gaps as its end-points, have $x$ fraction of loops inbetween, and then transition into a loop state of length $L$. Possible values of stochastic parameters are $l_1 \in [0,\infty)$, $l_2 \in [0,s]$, $h \in [0, s-l_2]$, $L \in [s-l_2-h, \infty]$ and $x \in [0,1]$. Expressions (4.3)-(4.6) fully define the linear part of the statistical properties of the chain, and will be used for the calculation of total contact probability.

### 5 Capture radius models

Building upon the framework of sections (2)-(4), we introduce a modification that takes into account the finite capture radius in Hi-C experiments, thus providing an explicit form of $\mathbb{P}(\text{contact}) := P(s)$ from Eq. (2.1). We do so in three distinct ways - via the most straightforward Gaussian kernel model, via the restriction fragments model, and via the simplified restriction fragments model. In all of the following sections we use the notation

$$\mathcal{P}(R \mid s) := \sum_{i=(a,b,c,d)} \langle P(R \mid \sigma_{(i)}) \rangle := \sum_{i=(a,b,c,d)} \int_{B_i} d\{A\}\; P(R \mid \sigma_{(i)}) \cdot \omega^{(i)}(\{A\}), \tag{5.1}$$

which corresponds to *averaging with respect to loop disorder*.

#### 5.1 Gaussian kernel model

The first way to define contact probability between two sites at contour distance $s$ in terms of spherical kernel $K(R)$, which bears information about contact radius $r_c$:

$$P(s) = 4\pi \int_0^\infty K(R)\mathcal{P}(R \mid s)R^2 dR, \tag{5.1.1}$$

where we have used spherical symmetry of distributions by integrating out angular variables. One notable way to define $K(R)$ is via Heaviside step function $K(R) = \Theta(R - r_c)$. In the limit $\sqrt{\langle R^2 \rangle} \gg r_c$ this case reproduces "loops, no cutoff" scenario that was considered in earlier works, and whose behaviour is depicted in Fig. 2B from the main text. Instead, here we will use $K(R)$ in the form of

$$K(R) = \exp\left(-\frac{3R^2}{2r_c^2}\right), \tag{5.1.2}$$

which smooths out contributions to contact probability from pairs with different spatial separations. While terms entering $\mathcal{P}(R \mid s)$ can take different forms, the underlying probability distribution function $P(R \mid \sigma)$ always has Gaussian form with some variance $\sigma^2$ and can be treated before averaging with respect to loop disorder, i.e. we can explicitly rewrite:

$$P(s) = \sum_{i=(a,b,c,d)} \int_{B_i} d\{A\}\, \omega^{(i)}(\{A\}) \underbrace{\int_0^\infty K(R)P(R \mid \sigma_i) \cdot 4\pi R^2 dR}_{z(r_c,\sigma_i)}, \tag{5.1.3}$$

where

$$P(R \mid \sigma_i) = \left(\frac{3}{2\pi\sigma_i^2}\right)^{3/2} \exp\left(-\frac{3R^2}{2\sigma_i^2}\right). \tag{5.1.4}$$

Thus, we can evaluate the highlighted part of Eq. (5.1.3), keeping in mind that the full contact probability consists of several such terms with various values of $\sigma_i$, but the same functional form. For the simplicity of expressions, in what follows we skip the index $i$ at $\sigma_i$. Calculating the highlighted part under the assumption of Gaussian kernel, we get

$$z(r_c, \sigma) = \frac{2^{3/2}\pi^{3/2} r_c^3 \sigma^3}{3^{3/2}(\sigma^2 + r_c^2)^{3/2}} \tag{5.1.5}$$

#### 5.2 Restriction fragments model

The second model we propose is grounded in the specific biochemical steps of the Hi-C protocol. After addition of a restriction enzyme, chromatin is cut into fragments of random length $v$. Because the enzyme cleaves whenever it encounters its recognition site—an event well approximated as a Poisson process—the resulting fragment-length distribution is exponential:

$$\rho_{\text{fragment}}(v) = \frac{1}{v_0} \exp\left(-v/v_0\right). \tag{5.2.1}$$

For two loci to be ligated into a chimeric product, the spatial volumes of their respective fragments (of lengths $v_1$ and $v_2$) must overlap. Consequently, a pair of loci separated by genomic distance $s$ can be captured as a contact only if their spatial separation $R$ is smaller than the effective combined size of the two fragments on which they reside:

$$R \leq c \cdot \sqrt{r^2(v_1) + r^2(v_2)}, \tag{5.2.2}$$

where $r(v) = bv^{1/2}$ is the spatial extent of a fragment of length $v$ and $c$ is a geometric factor reflecting the overlap criterion. Taking into account that fragment lengths are distributed exponentially, we arrive at

$$P(s) = \int_0^\infty \int_0^\infty \frac{dv_1 dv_2}{v_0^2} e^{-(v_1+v_2)/v_0} \int_0^{c \cdot b\sqrt{v_1+v_2}} \mathcal{P}(R \mid s) \cdot 4\pi R^2 dR, \tag{5.2.3}$$

where we have used spherical symmetry of distributions by integrating out angular variables. While, as in the previous section, terms entering $\mathcal{P}(R \mid s)$ can take different forms, the underlying probability distribution function $P(R \mid \sigma)$ always has Gaussian form with some variance $\sigma^2$, i.e. in the same manner as above we can explicitly rewrite:

$$P(s) = \sum_{i=(a,b,c,d)} \int_{B_i} d\{A\}\ \omega^{(i)}(\{A\}) \underbrace{\int_0^\infty \int_0^\infty \frac{dv_1 dv_2}{v_0^2} e^{-(v_1+v_2)/v_0} \int_0^{c \cdot b\sqrt{v_1+v_2}} P(R \mid \sigma_i) \cdot 4\pi R^2 dR}_{f(c,\sigma_i,v_0)}. \tag{5.2.4}$$

Thus, we can evaluate the highlighted part of Eq. (5.2.4), keeping in mind that the full contact probability consists of several such terms with various values of $\sigma_i$ (similarly, we will skip the index $i$ below), but the same functional form. Focusing on the rightmost integral, we obtain

$$\int_0^{c \cdot b\sqrt{v_1+v_2}} P(R \mid \sigma) \cdot 4\pi R^2 dR \sim \frac{2}{9}\pi\sigma^2 \left( \sqrt{6\pi}\sigma \cdot \mathrm{Erf}\left( \frac{\sqrt{\frac{3}{2}} bc\sqrt{v_1+v_2}}{\sigma} \right) - 6bc\sqrt{v_1+v_2}\, e^{-\frac{3b^2c^2(v_1+v_2)}{2\sigma^2}} \right). \tag{5.2.5}$$

Consequently, integrating expression (5.2.5) with double exponent in Eq. (5.2.4), we obtain

$$\int_0^\infty \int_0^\infty \frac{dv_1 dv_2}{v_0^2} e^{-(v_1+v_2)/v_0} (\dots) \implies f(c, \sigma, v_0) = \frac{2^{3/2} b^3 c^3 \pi^{3/2} v_0^{3/2} \sigma^3 (3b^2c^2v_0 + 5\sigma^2)}{(3b^2c^2v_0 + 2\sigma^2)^{5/2}}, \tag{5.2.6}$$

where $f(c, \sigma, v_0)$ is equal to the highlighted part of Eq. (5.2.4).

#### 5.3 One restriction fragment: a simplified model

We can consider a simplification of the model from the previous section by assuming that only one restriction fragment volume contributes to ligation, and by analogy with Eq.

(5.2.3), we obtain

$$P(s) = \int_0^\infty \frac{dv}{v_0} e^{-v/v_0} \int_0^{k \cdot b v^{1/2}} \mathcal{P}(R \mid s) \cdot 4\pi R^2 dR, \tag{5.3.1}$$

where $k$ plays the same function as $c$ in two-integral model, but can take a different numerical value. Making the same assumptions as in the previous section, we arrive at the result analogous to Eq. (5.2.6):

$$q(k, \sigma, v_0) = \frac{2^{3/2}\pi^{3/2}}{3^{3/2}} \cdot \frac{k^3 b^3 \sigma^3}{(k^2 b^2 + 2\sigma^2/3v_0)^{3/2}}. \tag{5.3.2}$$

.

#### 5.4 Equivalence of the capture models

We can notice that expressions (5.1.5), (5.2.6) and (5.3.2) can be mapped onto each other, thus making these models of capture formally equivalent. For example, we instantly see that the one-integral model (5.3.2) can be mapped onto the kernel model (5.1.5) by equating $q(k, \sigma, v_0) = z(r_c, \sigma)$ and obtaining

$$r_c^2 = 3k^2 b^2 v_0/2. \tag{5.4.1}$$

In order to connect the Gaussian kernel approach to the more accurate two-integral model from section (5.2), we can use the assumption that the contour distance $s$ between two loci is much larger than the typical restriction fragment length $v_0$, i.e. $s \gg v_0$. Since for typical cases $\sigma^2 = b^2 s$, this assumption translates into $\sigma^2 \gg b^2 v_0$, and we can reduce Eq. (5.2.6) to

$$f(c, v_0) \approx \frac{5}{2}\pi^{3/2} b^3 c^3 v_0^{3/2}, \tag{5.4.2}$$

thus removing dependency on $\sigma$. Using a similar argument, we can reduce Eq. (5.1.5) to

$$z(r_c) \approx \frac{2^{3/2}}{3^{3/2}}\pi^{3/2} r_c^3, \tag{5.4.3}$$

Equating it with Eq. (5.4.2), we get a relationship between the Gaussian kernel model and the two-fragments model

$$r_c^2 = \frac{3 \cdot 5^{2/3}}{2^{5/3}} \cdot c^2 b^2 v_0. \tag{5.4.4}$$

Doing the same to Eq. (5.3.2), one arrives to:

$$q(k, v_0) \approx \pi^{3/2} v_0^{3/2} k^3 b^3, \tag{5.4.5}$$

and, equating Eq. (5.4.5) with Eq. (5.4.2), we get the relationship between the two-fragments model and the simplified one

$$c^3 = \frac{2}{5} k^3, \tag{5.4.6}$$

thus making all the three models connected through the proper choice of the parameters $c, k$ and $r_c$. However, since we have no way of knowing specific values of coefficients $c$ and $k$ due to their inherently heuristic nature, we introduce the effective genomic unit

$$v_0^{\text{eff}} = k^2 v_0 \tag{5.4.7}$$

that takes into account geometric specifics of contact formation, as well as restriction-fragment statistics and cross-linking efficiency. Substituting it into the expression (5.4.1), we arrive at the relationship between the effective restriction fragment length and capture radius

$$r_c^2 = \frac{3}{2} b^2 v_0^{\text{eff}}, \tag{5.4.8}$$

thus providing equivalent ways to consider characteristic scales of contact formation in terms of either spatial or genomic distances.

### 6 Perturbation theory

We can calculate the resulting contact probability by directly integrating Eqs. (5.1) and (5.1.1) with contributions from Sections 3 and 4, which yields the red curve from Fig. 2B in the main text. However, it can be performed only numerically. Therefore, following the arguments provided in the main text, here we consider a small-scale perturbation of this model, focusing on ideal chain with fractal dimension $d_f = 2$ and contour distance $s \ll \lambda, g$, or, in other words, $s \ll T$, where $T = \lambda + g$ is the period of the chain. In order to understand which contributions to consider, we can rewrite statistical weights from Section 4 up to the linear precision, focusing on their subset represented in Figure S1B. Reducing Eqs. (4.3)-(4.5) to their simpler forms, we obtain the following approximations.

For diagram (a):

$$\omega^{(a)} = \frac{g}{g+\lambda} e^{-s/d} \approx \frac{g}{g+\lambda}\left(1 - \frac{s}{g}\right). \tag{6.1}$$

For diagram (b):

$$\omega^{(b)}(l, L) = \frac{1}{\lambda(g+\lambda)} \exp\left(-\frac{l}{g} - \frac{L}{\lambda}\right) \approx \frac{1}{\lambda(g+\lambda)} \exp\left(-\frac{L}{\lambda}\right) \cdot \left(1 - \frac{l}{g}\right), \tag{6.2}$$

where $L \in (0, \infty)$ and $l \in (Max[0, s-L], s)$. For diagram (c):

$$\omega^{(c)}(q, L) = \frac{L}{\lambda(g+\lambda)} \exp\left(-\frac{L}{\lambda}\right), \tag{6.3}$$

where $L \in (s, \infty)$ and $q \in (0, 1 - s/L)$. Performing integration of Eqs. (6.1), (6.2) and (6.3) with respect to stochastic parameters ($l$ and $L$ for diagram (b), $q$ and $L$ for diagram (c)), we obtain full probabilities $W^{(i)}$ of encountering the $i$-th diagram:

$$W^{(a)} \approx \frac{g}{g+\lambda} - \frac{s}{g} \cdot \frac{g}{g+\lambda} \tag{6.4}$$

$$W^{(b)} \approx \frac{s}{g+\lambda} \tag{6.5}$$

$$W^{(c)} \approx \frac{\lambda}{g+\lambda} - \frac{s}{g+\lambda} \tag{6.6}$$

From these expressions, we observe that

$$W^{(a)} + 2 * W^{(b)} + W^{(c)} \approx 1, \tag{6.7}$$

where 2 accounts for a symmetry factor. We see that, indeed, the considered configurations of the polymer chain constitute the linear order of expansion in the parameters $s/g$ and $s/\lambda$, and that diagrams from Figure S1B are sufficient.

As was shown in section (5.4), all capture models are equivalent, so we are going to use the form with the Gaussian kernel (5.1.1) for simplicity. Additionally, we are introducing the variable

$$\gamma(s) = b^2 s / r_c^2(v_0^{\text{eff}}) = 2s/3v_0^{\text{eff}}, \tag{6.8}$$

which corresponds to genomic distance between contacts in terms of effective restriction fragment units (in what follows, we omit the dependence on $s$ from notation, writing $\gamma$ instead of $\gamma(s)$). Expanding the expression for contact probability $P(s)$, we obtain

$$P(s) = 4\pi \int_0^\infty K(R) R^2 dR \Big( W^{(a)} P^{(a)}(R \mid s) + 2 \cdot \int_0^\infty dL \int_{Max[s-L,0]}^s \omega^{(b)}(l, L) P^{(b)}(R \mid s, l, L) dl + \\ + \int_s^\infty dL \int_0^{1-s/L} \omega^{(c)}(q, L) P^{(c)}(R \mid s, L) dq \Big). \tag{6.9}$$

For the first term in Eq. (6.9), which corresponds to diagram (a), we get:

$$\frac{4\pi}{(2\pi b^2 s/3)^{3/2}} \cdot \frac{g}{g+\lambda} \cdot \left(1 - \frac{s}{g}\right) \int_0^\infty K(R) R^2 \exp\left(-\frac{3R^2}{2b^2 s}\right) dR. \tag{6.10}$$

For the second term, which corresponds to diagram (b), we get:

$$\frac{8 \cdot 3^{3/2} \cdot \pi s^{1/2}}{(2\pi b^2)^{3/2} \cdot \lambda \cdot (g+\lambda)} \int_0^\infty K(R) R^2 dR \int_0^1 d\tilde{L} \int_{1-\tilde{L}}^1 d\tilde{l} \frac{\tilde{L}^{3/2} e^{-\tilde{L}s/\lambda} \cdot (1 - \tilde{l}s/g)}{(\tilde{L} + 2\tilde{l} - \tilde{l}^2 - 1)^{3/2}} \exp\left(-\frac{3R^2}{2b^2 s} \frac{\tilde{L}}{\tilde{L} + 2\tilde{l} - \tilde{l}^2 - 1}\right) + \\ + \frac{8 \cdot 3^{3/2} \cdot \pi s^{1/2}}{(2\pi b^2)^{3/2} \cdot \lambda \cdot (g+\lambda)} \int_0^\infty K(R) R^2 dR \int_1^\infty d\tilde{L} \int_0^1 d\tilde{l} \frac{\tilde{L}^{3/2} e^{-\tilde{L}s/\lambda} \cdot (1 - \tilde{l}s/g)}{(\tilde{L} + 2\tilde{l} - \tilde{l}^2 - 1)^{3/2}} \exp\left(-\frac{3R^2}{2b^2 s} \frac{\tilde{L}}{\tilde{L} + 2\tilde{l} - \tilde{l}^2 - 1}\right) \tag{6.11}$$

And, finally, for the third term, which corresponds to diagram (c), we get:

$$\frac{4\pi s^{1/2}}{(2\pi b^2/3)^{3/2} \cdot \lambda(g+\lambda)} \int_0^\infty K(R) R^2 dR \int_1^\infty d\tilde{L} \frac{\tilde{L}^{3/2} e^{-\tilde{L}s/\lambda}}{(\tilde{L}-1)^{1/2}} \exp\left(-\frac{3R^2}{2b^2 s} \frac{\tilde{L}}{\tilde{L}-1}\right). \tag{6.12}$$

Substituting the specific form of $K(R)$ given by Eq. (5.1.2), for diagram (a), represented by

Eq. (6.10), we obtain

$$\frac{4\pi}{(2\pi b^2 s/3)^{3/2}} \cdot \frac{g}{g+\lambda} \cdot \left(1-\frac{s}{g}\right) \cdot \frac{\sqrt{\pi/2}}{3^{3/2}\cdot(1/r_c^2(v)+1/b^2 s)^{3/2}} = \frac{1}{(\gamma+1)^{3/2}} \cdot \frac{g-s}{g+\lambda}. \tag{6.13}$$

For diagram (c), represented by Eq. (6.12), we get

$$\frac{4\pi s^{1/2}}{(2\pi b^2)^{3/2}\cdot\lambda(g+\lambda)} \int_1^\infty d\tilde{L} \frac{\tilde{L}^{3/2} e^{-\tilde{L}s/\lambda}}{(\tilde{L}-1)^{1/2}} \cdot \frac{\sqrt{\pi/2}}{\left(\frac{1}{r_c^2(v)}+\frac{\tilde{L}}{b^2 s(\tilde{L}-1)}\right)^{3/2}} = \frac{s^2}{\lambda(g+\lambda)} \int_1^\infty \frac{(\tilde{L}-1)e^{-\tilde{L}s/\lambda}}{\left(1+\gamma\left(\frac{\tilde{L}-1}{\tilde{L}}\right)\right)^{3/2}} d\tilde{L} \tag{6.14}$$

We have to asymptotically expand this expression in terms of parameter $s/\lambda \overset{\text{def}}{=} \alpha \ll 1$. Performing integration by parts to separate divergent contributions, we obtain

$$\begin{aligned}
&\int_1^\infty \frac{(\tilde{L}-1)e^{-\tilde{L}\alpha}}{\left(1+\gamma\left(\frac{\tilde{L}-1}{\tilde{L}}\right)\right)^{3/2}} d\tilde{L} = \\
&= \left(-\frac{(\tilde{L}-1)}{\left(1+\gamma\left(\frac{\tilde{L}-1}{\tilde{L}}\right)\right)^{3/2}} \cdot \frac{e^{-\tilde{L}\alpha}}{\alpha}\right)\Bigg|_{\tilde{L}=1}^{\tilde{L}=\infty} - \left(\frac{3\gamma+2(\gamma+1)\tilde{L}^2-5\gamma\tilde{L}}{2(\gamma(\tilde{L}-1)+\tilde{L})^2\sqrt{\gamma-\frac{\gamma}{\tilde{L}}+1}} \frac{e^{-\tilde{L}\alpha}}{\alpha^2}\right)\Bigg|_{\tilde{L}=1}^{\tilde{L}=\infty} + \\
&+ \int_1^\infty \frac{3\gamma\left(\gamma(\tilde{L}-1)-4\tilde{L}\right)}{4\sqrt{\tilde{L}}(\gamma(\tilde{L}-1)+\tilde{L})^{7/2}} \frac{e^{-\tilde{L}\alpha}}{\alpha^2} d\tilde{L} = \\
&= \frac{e^{-\alpha}}{\alpha^2} + \frac{1}{\alpha^2} I(\alpha),
\end{aligned} \tag{6.15}$$

where

$$I(\alpha) = \int_1^\infty \frac{3\gamma\left(\gamma(\tilde{L}-1)-4\tilde{L}\right)}{4\sqrt{\tilde{L}}(\gamma(\tilde{L}-1)+\tilde{L})^{7/2}} e^{-\tilde{L}\alpha} d\tilde{L} \tag{6.16}$$

is a convergent integral that can be calculated by Taylor expanding $e^{-\tilde{L}\alpha}$:

$$I(\alpha) \approx \frac{1}{(\gamma+1)^{3/2}} - 1 - \alpha\left(-\frac{\gamma}{2(\gamma+1)^{5/2}} + \frac{1}{(\gamma+1)^{5/2}} - 1\right) \tag{6.17}$$

Final contribution from diagram (c) is thus given by

$$\frac{s^2}{\lambda(g+\lambda)} \cdot \left(\frac{e^{-\alpha}}{\alpha^2} + \frac{1}{\alpha^2} \cdot I(\alpha)\right) \approx \frac{\lambda}{g+\lambda} \cdot \left(\frac{1}{(\gamma+1)^{3/2}} + \frac{s}{\lambda} \cdot \frac{\gamma-2}{2(\gamma+1)^{5/2}}\right) \tag{6.18}$$

For diagram (b), represented by Eq. (6.11), we get:

$$\frac{8\pi s^{1/2}}{(2\pi b^2)^{3/2}\cdot\lambda\cdot(g+\lambda)}\cdot\left(\int_0^1 d\tilde{L}\int_{1-\tilde{L}}^1 d\tilde{l}+\int_1^\infty d\tilde{L}\int_0^1 d\tilde{l}\right)\frac{\tilde{L}^{3/2}e^{-\tilde{L}s/\lambda}\cdot(1-\tilde{l}s/g)}{(\tilde{L}+2\tilde{l}-\tilde{l}^2-1)^{3/2}}\frac{\sqrt{\pi/2}}{\left(\frac{1}{r_c^2}+\frac{\tilde{L}}{b^2 s\left(\tilde{L}-(\tilde{l}-1)^2\right)}\right)^{3/2}}=$$

$$=\frac{2s^2}{\lambda(g+\lambda)}\left(\int_0^1 d\tilde{L}\int_{1-\tilde{L}}^1 d\tilde{l}+\int_1^\infty d\tilde{L}\int_0^1 d\tilde{l}\right)\frac{\left(1-\tilde{l}s/g\right)e^{-\tilde{L}s/\lambda}}{\left(\gamma\left(1-\frac{(\tilde{l}-1)^2}{\tilde{L}}\right)+1\right)^{3/2}}=$$

$$=\frac{2s^2}{\lambda(g+\lambda)}\int_0^1 dL\frac{Le^{-\frac{Ls}{\lambda}}\left(\gamma\sqrt{\gamma+1}g+s\left(\sqrt{\gamma+1}-\gamma\sqrt{\gamma-\gamma L+1}-\sqrt{\gamma-\gamma L+1}\right)\right)}{\gamma(\gamma+1)^{3/2}g\sqrt{\gamma-\gamma L+1}}+$$

$$+\frac{2s^2}{\lambda(g+\lambda)}\int_1^\infty dLe^{-\frac{Ls}{\lambda}}\left(\frac{s\left(\sqrt{(\gamma+1)L(-\gamma+\gamma L+L)}-(\gamma+1)L\right)}{\gamma(\gamma+1)^{3/2}g}+\frac{1}{(\gamma+1)\sqrt{\gamma-\frac{\gamma}{L}+1}}\right) \tag{6.19}$$

It is easy to check that the former integral in this sum doesn't give a linear contribution due to finite integration limits, so the whole contribution is defined by the second term:

$$\frac{2s^2}{\lambda(g+\lambda)(\gamma+1)}\int_1^\infty dLe^{-L\alpha}\left(\frac{1}{\sqrt{\gamma-\gamma/L+1}}+\frac{1}{\gamma}\frac{s}{d}\left(\sqrt{L(\gamma(L-1)+L)}-(\gamma+1)^{1/2}L\right)\right) \tag{6.20}$$

This expression has separate terms, which we will consecutively analyse. For the first one, we get

$$\begin{aligned}\int_1^\infty dL\frac{e^{-L\alpha}}{\sqrt{\gamma-\gamma/L+1}}&=\left(-\frac{1}{\sqrt{\gamma-\gamma/L+1}}\cdot\frac{e^{-\tilde{L}\alpha}}{\alpha}\right)\Bigg|_{\tilde{L}=1}^{\tilde{L}=\infty}-\\ -\int_1^\infty\frac{\gamma}{2\sqrt{L}(\gamma(L-1)+L)^{3/2}}\frac{e^{-L\alpha}}{\alpha}&=\frac{e^{-a}}{a}-\frac{1}{\alpha}I(\alpha)\end{aligned} \tag{6.21}$$

$$I(\alpha)\approx 1-\frac{1}{\sqrt{\gamma+1}} \tag{6.22}$$

Final contribution from the first term:

$$\frac{e^{-\alpha}}{\alpha}-\frac{1}{\alpha}\left(1-\frac{1}{\sqrt{\gamma+1}}\right)\approx\frac{1}{\alpha\sqrt{\gamma+1}}+\frac{\alpha}{2}-1 \tag{6.23}$$

For the second term:

$$\begin{aligned}&\int_1^\infty dLe^{-L\alpha}\sqrt{L(\gamma(L-1)+L)}=\left(-\sqrt{L(\gamma(L-1)+L)}\cdot\frac{e^{-\tilde{L}\alpha}}{\alpha}\right)\Bigg|_{\tilde{L}=1}^{\tilde{L}=\infty}-\\ &-\left(\frac{(\gamma+1)L+\gamma(L-1)+L}{2\sqrt{L(\gamma(L-1)+L)}}\frac{e^{-\tilde{L}\alpha}}{\alpha^2}\right)\Bigg|_{\tilde{L}=1}^{\tilde{L}=\infty}-\frac{1}{\alpha^2}\int_1^\infty dL\frac{\gamma^2}{4(L(\gamma(L-1)+L))^{3/2}}e^{-L\alpha}=\\ &=\frac{e^{-\alpha}}{\alpha}+\frac{e^{-\alpha}(\gamma+2)}{2\alpha^2}-\frac{1}{\alpha^2}I(\alpha)\end{aligned} \tag{6.24}$$

$$I(\alpha) \approx \frac{1}{2}\left(\gamma - 2\sqrt{\gamma+1} + 2\right) \tag{6.25}$$

Final contribution from the second term:

$$\frac{e^{-\alpha}}{\alpha} + \frac{e^{-\alpha}(\gamma+2)}{2r_c^2} - \frac{1}{2r_c^2}\left(\gamma - 2\sqrt{\gamma+1} + 2\right) \approx \frac{\sqrt{\gamma+1}}{r_c^2} + \frac{1}{12}\alpha(4-\gamma) - \frac{\gamma}{2\alpha} + \frac{\gamma-2}{4} \tag{6.26}$$

And, finally, for the third term:

$$\int_1^\infty dL L e^{-L\alpha} = \frac{(\alpha+1)e^{-\alpha}}{r_c^2} \approx \frac{1}{r_c^2} + \frac{\alpha}{3} - \frac{1}{2} \tag{6.27}$$

Substituting these terms into Eq. (6.19) and considering only linear contributions, we see that Eq. (6.23) yields

$$\frac{2s}{(g+\lambda)(\gamma+1)^{3/2}} \tag{6.28}$$

Eq. (6.26) yields

$$\frac{2s}{g+\lambda}\frac{\lambda}{g}\frac{1}{\gamma(\gamma+1)^{1/2}} \tag{6.29}$$

And Eq. (6.27) yields

$$-\frac{2s}{g+\lambda}\frac{\lambda}{g}\frac{1}{\gamma(\gamma+1)^{1/2}} \tag{6.30}$$

Summing these three contributions with Eqs (6.13) and (6.18) we obtain the final answer for contact probability, where we remind that we put $\gamma = b^2 s / r_c^2(v_0^{\text{eff}}) = 2s/3v_0^{\text{eff}}$:

$$\boxed{P(s) = \frac{1}{(\gamma+1)^{3/2}}\left(1 + \frac{s}{g+\lambda}\frac{3\gamma}{2(\gamma+1)}\right).} \tag{6.31}$$

Logarithmic derivative of this expression is given by

$$y(s) = \frac{d\log_{10}P(s)}{d\log_{10}s} = \frac{s(3\gamma(\gamma+2))}{(g+\lambda)\left(2(\gamma+1)^2\right)} - \frac{3\gamma}{2(\gamma+1)} \tag{6.32}$$

It has a minimum that we can find by computing a second-order derivative

$$G(s) = \frac{d\,y(s)}{ds} = -\frac{3\gamma(\gamma g + g + \lambda + \gamma\lambda - s(\gamma(\gamma+3)+4))}{2s(g+\lambda)(\gamma+1)^3} \tag{6.33}$$

Assuming $\gamma \gg 1$, we find that

$$\frac{2(g+\lambda)}{3}G(s) = -\frac{\gamma(\lambda+g)}{s(\gamma+1)^2} + \frac{4\gamma}{(\gamma+1)^3} + \frac{3\gamma^2}{(\gamma+1)^3} + \frac{\gamma^3}{(\gamma+1)^3} \approx 1 - \frac{1}{s}\frac{\lambda+g}{\gamma} \tag{6.34}$$

Therefore, the minimum of Eq. (6.32) is approximately located at

$$s_{\min} = \sqrt{r_c^2(v_0^{\text{eff}})\cdot(g+\lambda)/b^2} = \sqrt{3/2\cdot v_0^{\text{eff}}\cdot(g+\lambda)}, \tag{6.35}$$

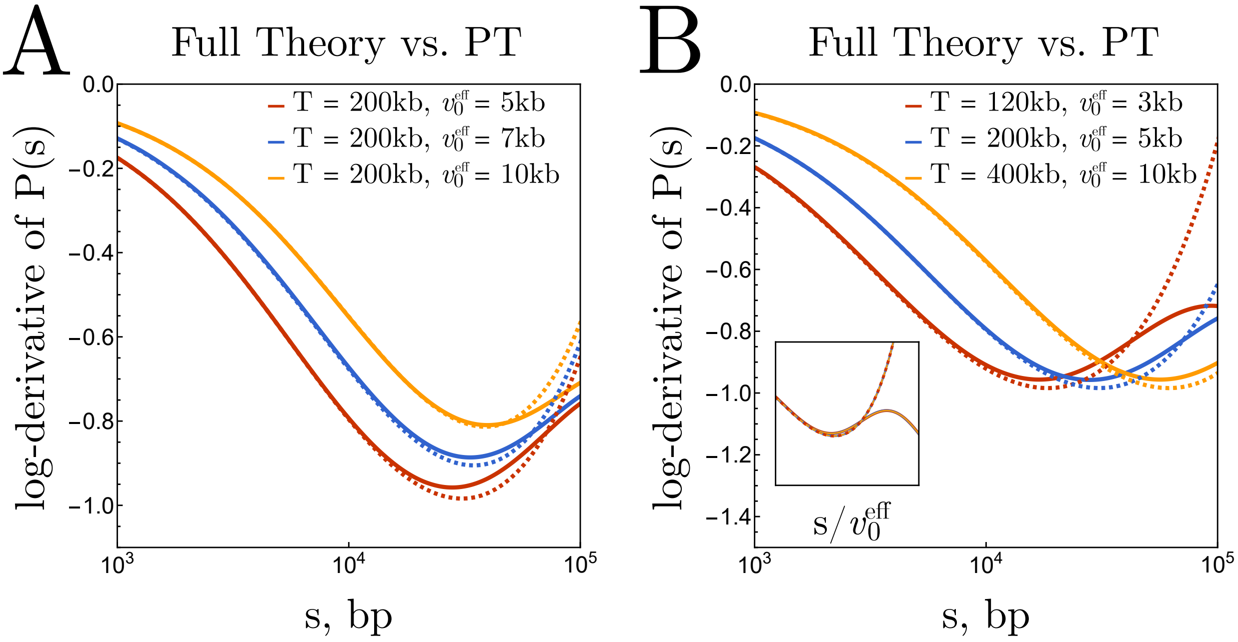

Figure S2: Comparison of full theory (solid lines) and perturbation theory (dashed lines) predictions. (A) Logarithmic derivative of $P(s)$ for fixed loop–gap period $T$ and varying restriction fragment lengths $v_0^{\text{eff}}$. For all curves, $\lambda = 130$kb and $g = 70$kb. (B) Logarithmic derivative of $P(s)$ for constant ratio $T/v_0^{\text{eff}} = 40$. Red curve: $\lambda = 90$kb, $g = 30$kb. Blue curve: $\lambda = 120$kb, $g = 80$kb. Orange curve: $\lambda = 250$kb, $g = 150$kb. Inset: rescaling $s$ by $v_0^{\text{eff}}$ leads to a collapse.

where the value of logarithmic derivative is approximately given by

$$y_{\min} \approx \frac{6 - 3(\frac{\gamma}{s} s_0)^2}{2(1 + \frac{\gamma}{s} s_0)^2}. \tag{6.36}$$

We note that Eqs. (6.35) and (6.36) require the assumption that $s \gg v_0^{\text{eff}}$, in addition to the perturbative assumption $s \ll T$. Therefore, these simple expressions show some degree of discrepancy with the position of the dip obtained directly from Eq. (6.31) (see Fig. S3). Nevertheless, they manage to capture the qualitative behaviour of the model and can be used for approximate inferences, whereas quantitative evaluations should be performed using Eqs. (6.31) and (6.32) instead.

### 7 MD simulations

To test the predictions of our analytical framework and to investigate 3D folding of looped polymers, we performed 3D molecular dynamics simulations of the chains combining 1D *active loop extrusion* on top of two different equilibrated systems: phantom and crumpled chains (see below). Each cohesin from 1D trajectory was represented as an additional harmonic bond between two beads. 3D polymer simulations are done using Polychrom, a wrapper around the open-source GPU-assisted molecular dynamics package OpenMM (Eastman and Pande, 2010). At the end of simulation, we computed steady-state contact probabilities and compared them

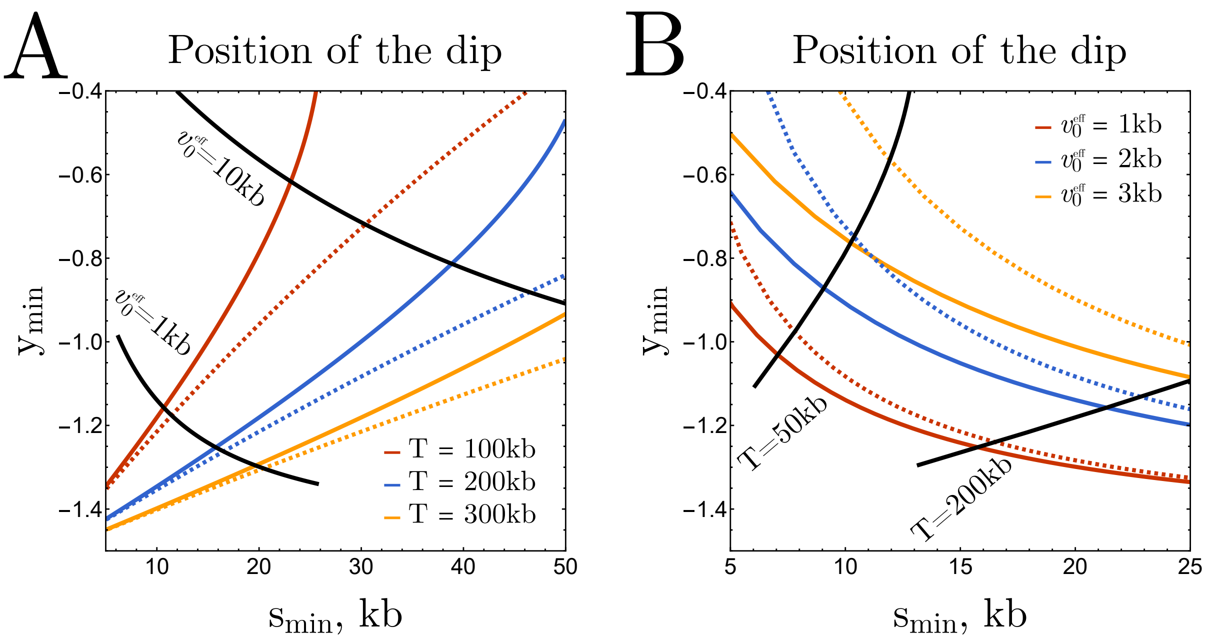

Figure S3: Discrepancies between position of the dip obtained numerically from the perturbation theory expression (6.31) (solid lines), and its position obtained analytically under the additional limit $s \gg v_0^{\rm eff}$, given by Eqs (6.35) and (6.36) (dashed lines). (A) Constant period $T$ and variable restriction fragment length $v_0^{\rm eff}$ along each colored curve. (B) Constant restriction fragment size $v_0^{\rm eff}$ and variable period $T$ along each colored curve.

against perturbation theory (PT).

**1D extrusion**. In the 1D simulations of extrusion, cohesins followed the state-of-the-art extrusion kinetics, characterized by three parameters: the mean cohesin spacing $d$ (inverse linear density), cohesin processivity $l$ and extrusion rate $r$. At initialization, $N_0 = N/d$ cohesins were placed randomly along the chain, defining the mean spacing $d$. During simulations, each bound cohesin unbound with rate $k_{\rm off} = 2r/l$, where $l$ is the processivity length and $r$ the extrusion speed, and the same number of cohesins were reloaded at random unoccupied sites to maintain $N_0$ constant. Upon binding, cohesins extruded symmetrically at speed $r \approx 1$ kb/s until unbinding, collision with another cohesin, or reaching the chain ends. Simulation time between extrusion steps was mapped to real time ($0.8{\rm kb}/r = 0.8$s) by calibrating the Rouse diffusion coefficient of individual beads, yielding $D_R \approx 10^{-2}\,\mu{\rm m}^2\,{\rm s}^{-1/2}$, consistent with experimental measurements (see Gabriele et al, 2022; Brückner et al, 2023).

**Phantom chains**. To benchmark perturbation theory in the simplest case, we first simulated phantom chains with random-walk statistics, i.e. as phantom beads ($N = 40000$) on harmonic springs with excluded-volume interactions and angular potential switched off. The only potential in the model in this case was the harmonic nearest-neighbor potential:

$$U_{\rm bond} = \frac{3}{2a^2}\sum_{i=1}^{N-1}(r_{i,i+1} - l_b)^2, \tag{7.1}$$

where $a = 0.06\sigma$ is the standard deviation of the monomer-to-monomer distance $r_{i,i+1} = |\mathbf{r}_{i+1} - \mathbf{r}_i|$ from the equilibrium bond length $l_b = \sigma$.

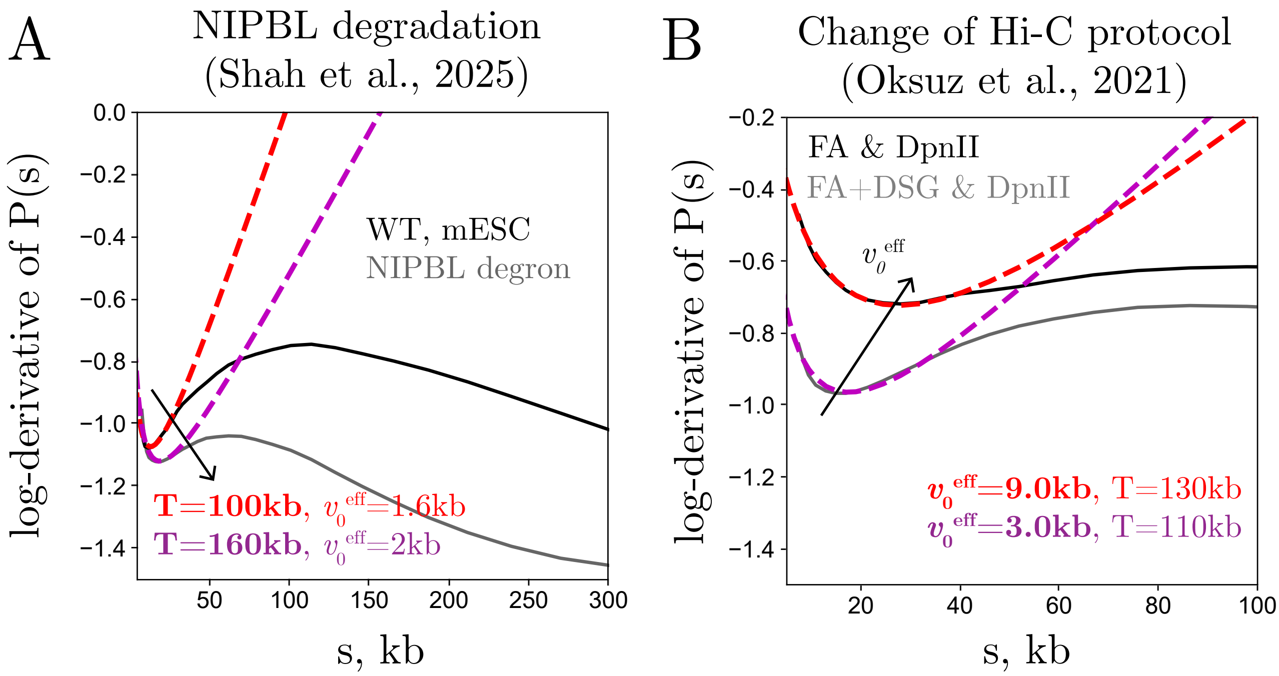

Figure S4: Experimental validation of the theoretical framework. (A) Degradation of NIPBL (cohesin loader) in mESCs leads to a consistent decrease in loop density from ∼10 to ∼6.25 loops/Mb, while $v_0^{\text{eff}}$ remains almost unchanged, as predicted by the theory. Colored curves represent best-fit theoretical profiles with parameters indicated in the legend. Hi-C data from Shah et al. (B) Improved cross-linking enhances Hi-C resolution. Comparison of two protocols in human embryonic stem cells (ESC cells) from Oksuz et al: standard in situ Hi-C (FA crosslinking, DpnII digestion) versus enhanced Hi-C (FA+DSG crosslinking, DpnII digestion). Red/magenta theory fits show a threefold decrease in $v_0^{\text{eff}}$ upon addition of DSG, while loop density remains nearly constant.

**Crumpled chains**. In the absence of cohesin, Hi-C and Micro-C experiments show $P(s) \sim s^{-\gamma_c}$ with $\gamma_c \approx 1$, consistent with a crumpled polymer of fractal dimension $d_f \approx 3$ (Polovnikov et al, PRX 2023). The canonical polymer model that reproduces this behavior is an equilibrium melt of unknotted, non-concatenated ring polymers, where each polymer chain exhibits a crumpled regime with $d_f = 3$ beyond the entanglement length $s > N_e$, i.e. the end-to-end spatial distance $R(s)$ of a segment of contour length $s$ has the following behaviour:

$$R(s) = \begin{cases} b\, s^{1/2}, & s < N_e \\ b\, s^{1/3} N_e^{1/6}, & s > N_e. \end{cases} \tag{7.2}$$

The entanglement length $N_e$ defines a crossover length scale from ideal ($d_f = 2$) to crumpled ($d_f = 3$) folding. Using the mean-field argument, $P(s) \sim R(s)^{-1/3}$, one can find that the experimentally observed power-law $P(s) \sim s^{-1}$ is feature of the crumpled state with $d_f = 3$.

Aimed at realistic chromatin baseline without loops, we thus simulated a single unknotted ring polymer of size $N = 40000$ in periodic boundary conditions. The chain was equipped with harmonic bonds $U_{\text{bond}}$ (Eq. (7.1)), quadratic angular potential $U_{\text{angle}}$ and excluded volume $U_{\text{ev}}$ interactions. The excluded volume potential $U_{\text{ev}}$ was introduced via the auxiliary Weeks-Chandler-Anderson (WCA) potential $U(r_i, r_j)$, which is a lifted Lennard-Jones repulsive branch

$$U(r_{ij} = |\mathbf{r_i} - \mathbf{r_j}|) = \begin{cases} 4\varepsilon\left((\sigma/r_{ij})^{12} - (\sigma/r_{ij})^{6}\right) + \varepsilon, & r_{ij} \le 2^{1/6}\sigma \\ 0, & r_{ij} > 2^{1/6}\sigma \end{cases} \tag{7.3}$$

where $\sigma$ is the characteristic scale of the excluded volume repulsion and $\varepsilon = 1$. In order to avoid strong repulsive forces in Eq. (7.3) at extremely short distances $r_{ij} \ll \sigma$, the WCA potential was further smoothly truncated as follows:

$$U_{\text{ev}}(r_{ij}) = \mathcal{H}(U(r_{ij}) - \varepsilon_{tr})\varepsilon_{tr} \times \left(1 + \tanh\left[\frac{U(r_{ij})}{\varepsilon_{tr}} - 1\right]\right) + \mathcal{H}(\varepsilon_{tr} - U(r_{ij}))U(r_{ij}), \tag{7.4}$$

at the prescribed truncation value $\varepsilon_{tr} = 10$, corresponding to a strong mutual volume exclusion of the beads. In Eq. (7.4) $\mathcal{H}$ is the step function. The potential Eq. (7.4) acts between every pair of beads, except for neighboring ones. We further imposed the bond length slightly smaller than the scale of excluded volume interaction in this simulation ($l_b = 0.8\sigma$) to ensure the bonds do not cross in the course of simulation.

The angular energy $U_{\text{angle}}$ was introduced as a harmonic quadratic potential for the angle $\theta_{i,i+1,i+2}$ between the two consecutive bonds

$$U_{\text{angle}} = \sum_{i=1}^{N-2} \frac{1}{2} k(\theta_{i,i+1,i+2} - \theta_0)^2 \tag{7.5}$$

with parameters $k = 1$ and $\theta_0 = \pi$.

Simulations were performed at a polymer volume density of $\rho\sigma^3 = 0.4$, controlled by adjusting the simulation box size. This density most accurately reproduced Micro-C data from mouse embryonic stem cells under cohesin-depleted conditions (Fig. S5D, S7B). By fitting the simulated log-derivative of $P(s)$ to the experimental Micro-C profiles, we also inferred an optimal chromatin compaction of 800 bp per bead. Using this mapping, the effective persistence length of the model was estimated as $l_p \approx 1.8$ kb via comparison with a worm-like chain benchmark (Fig. S7A), consistent with experimental measurements in yeast (Arbona *et al.*, 2017). The associated entanglement length was found to be around $N_e \approx 100$ kb.

This calibration of the loop-free baseline model against Micro-C data over genomic separations from $s = 5$ kb to $s \sim 10$ Mb provides an optimized foundation for loop-free chromatin simulations. Although such full-scale calibration may appear unnecessary for the present study—focused primarily on short genomic distances $s < N_e$ where ideal statistics dominate (see Eq.(7.2))—it appears as a by-product of calibration of the volume density, affecting the persistence length $l_p$ and the short scale statistics.

Interestingly, Micro-C data from cohesin-depleted mESC cells show a wedge-shaped log-derivative profile (Fig. S5D, S7B), rather than the generally expected plateau seen in Hi-C under similar cohesin-free conditions. We demonstrated that this wedge arises naturally when the fragment length $v_0$ used in simulations is smaller than the chromatin persistence length $l_p$, in agreement with the persistence correction (see also below in Eq. (7.6)). The fragment length used to recapitulate the Micro-C curve ($v_0 \approx 0.8$ kb) is notably less than $l_p = 1.8$ kb, explaining the wedge (Fig. S7B). By contrast, the Hi-C datasets without cohesin display the familiar plateau-like profiles at scales $\approx$ 30kb-3Mb, which our simulations also reproduce with larger $v_0$ ($> l_p$) (Fig. S7B). Therefore, varying only $v_0$ reconciles various Hi-C and Micro-C datasets without cohesin, revealing a universal loops-free baseline of chromosome organization.

**Computing the contact probability $P(s)$ in simulations**. After equilibration of both (phantom and crumpled) systems with or without loops on top, simulation snapshots were collected and contact probabilities $P(s)$ were computed as follows: bead–bead contacts were registered at capture distances $r_c = R(v)$ corresponding to the typical end-to-end spatial distance $R(v)$ of polymer segments of various contour length $v$. The raw contact counts $\mathcal{N}(R(v))$ were then weighted by the exponential fragment-length distribution $\rho(v|v_0) = v^{-1}\exp(-v/v_0)$, simulating the fragmentation protocol in Hi-C. This yielded the contact probability for each $v_0$ of the following form:

$$P(s|v_0) = \sum_{i=1}^{k} \mathcal{N}(R(v_i))\, \rho(v_i|v_0)\Delta v,$$

where $k = 20$ fragment sizes were used, equally spaced between $v_1 = 1$ kb and $v_k = 4v_0$, with step size $\Delta v = (v_k - v_1)/k$. This approach is numerically equivalent to the analytical fragment model described in Section 5.3 with the actual spatial size of the fragments $R(v)$ computed directly in simulations rather than using the ideal-chain benchmark $k\, bv^{1/2}$.

The key difference between the theoretical model and the crumpled-chain simulations

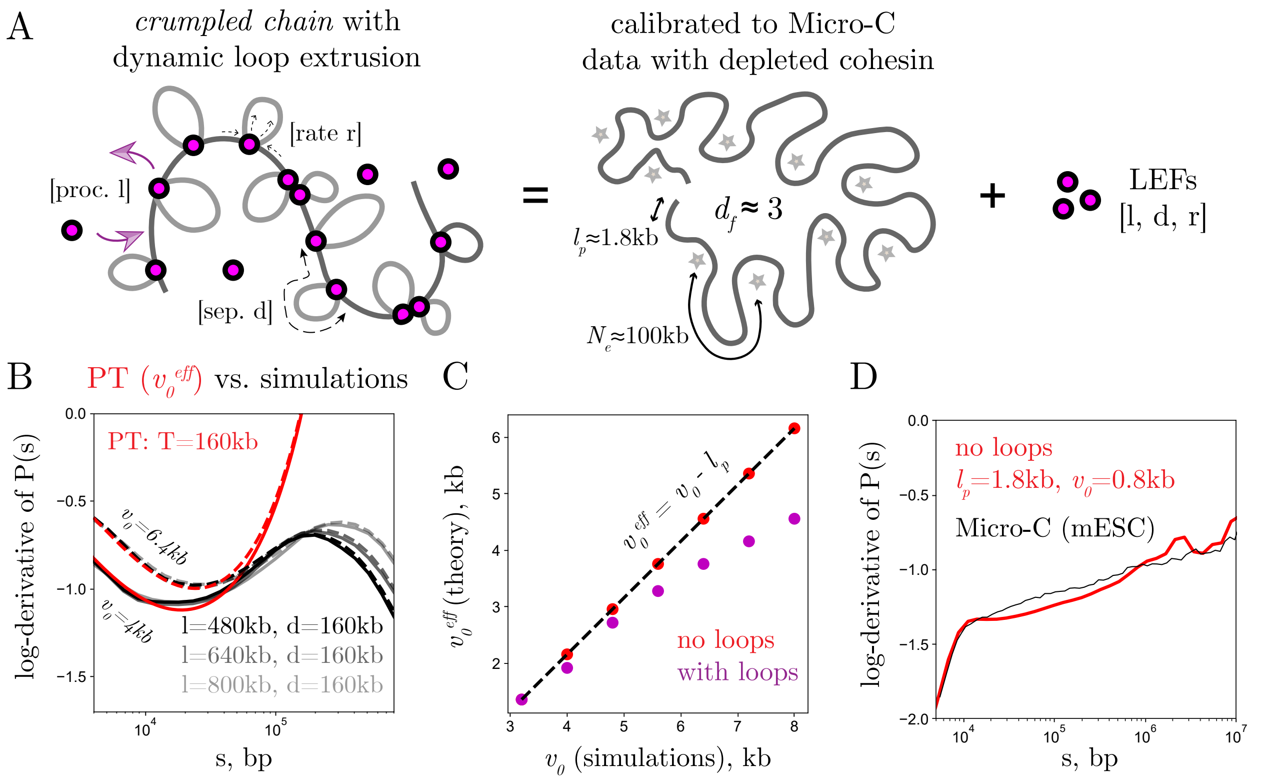

Figure S5: (A) Schematic of a crumpled polymer chain, calibrated against Micro-C data, with active loop extruders. (B) Theory-simulation comparison for crumpled chains using $T = d$ and the effective fragment length parameter in theory $v_0^{\text{eff}} = v_0 - l_p$, where $v_0$ is the mean fragment length in simulations; two curves are shown for $v_0 = 4$ kb (solid; $v_0^{\text{eff}} = 2.2$kb) and $v_0 = 6.4$ kb (dashed; $v_0^{\text{eff}} = 4.6$kb). (C) Relationship between $v_0$ in simulations and $v_0^{\text{eff}}$ values in PT for both looped and unlooped systems. (D) Calibration of loops-free crumpled chain simulation against Micro-C data from Hsieh et al. for cells without cohesin. Curve matching at $s < 10$ kb enables unit conversion between simulations and biological chromatin (800bp per bead, the compaction ratio assumed for both phantom and crumpled chain simulations) and persistence length $l_p = 1.8$kb.

is that, in the latter (but not in the phantom-chain case), restriction fragments are semi-flexible and therefore deviate from the ideal-chain benchmark. We therefore sought to determine how this intrinsic persistence modifies the relation between the theoretical parameter $v_0^{\text{eff}}$—the effective fragment length entering the Gaussian kernel model defined in Eq. (5.4.8)—and the actual genomic length of semi-flexible restriction fragments $v_0$.

For phantom random-walk chains, the situation is straightforward: the effective fragment length equals the true fragment length, $v_0^{\text{eff}} = v_0$, as follows directly from Eqs. (5.3.1) and (5.4.7), in full agreement with simulation results. In contrast, for semi-flexible crumpled chains, the intrinsic persistence suppresses short-range contact formation within fragments smaller than (or comparable to) the persistence length, thereby reducing the effective capture radius relative to the ideal-chain prediction. As shown below, this leads to a systematic reduction of $v_0^{\text{eff}}$ by approximately one persistence length $l_p$, a correction that reconciles the theoretical flexible-chain model with the simulation data for semi-flexible polymers.

**Results for crumpled-chain simulations with loops**. Building on the calibrated crumpled-chain model, we next introduced active loop extrusion following the procedure described in the corresponding paragraph above. Then we compared the resulting contact curves with theoretical predictions (perturbation theory, PT) for flexible chains with quenched loops. Remarkably, as in the phantom-chain case (Fig. S6), PT again reproduced the simulated log-derivatives, provided simulated fragment length $v_0$ is reduced by $l_p$ for the effective fragment length $v_0^{\text{eff}}$ in PT: $v_0^{\text{eff}} = v_0 - l_p$, where $l_p = 1.8$kb is the chromatin persistence length (Fig. S5C,D).

This correction reflects the intrinsic semi-flexibility of chromatin: restriction fragments shorter than (or comparable to) $l_p$ behave as nearly rigid, impermeable blobs, so for $v_0 \approx l_p$ very few short-range contacts can be registered and effective fragment length $v_0^{\text{eff}}$ (and capture radius $r_c$) is practically zero. Conversely, for large fragments $v_0 \gg l_p$, the chain behaves effectively flexible on the relevant scales, and PT agrees directly with simulations ($v_0^{\text{eff}} \simeq v_0$), consistent with the scale invariance of the flexible-chain log-derivative under simultaneous rescaling of all lengths $(s, v_0^{\text{eff}}, T)$ by $l_p$. These considerations motivate the simple ansatz

$$v_0^{\text{eff}} = v_0 - l_p, \qquad v_0 \geq l_p, \tag{7.6}$$

which quantitatively reconciles crumpled-chain simulations with the theoretical model. Equation (7.6) also explains the disappearance of the dip when $v_0 = l_p$: at this point the effective fragment length reduces to the bead size $b$ of a fully flexible chain of persistence blobs, corresponding to a vanishing effective capture radius (blue theoretical curve in Fig. 2B).

We first validated Eq. (7.6) in crumpled chains without loops (Fig. S5C, Fig. S8A). In this case, log-derivatives are monotonic and lack the dip, as expected from the theory (orange curve in Fig. 2B). Using only the zeroth order in PT and (7.6), we matched the entire family of simulation curves (Fig. S8A) across $v_0$, confirming the persistence correction.

Introducing loops of density $d^{-1}$, we again found fair agreement at $v_0$ across the reasonable lengths of restriction fragments obtained in Hi-C with typical enzymes (up to 5–6 kb;

e.g., DpnII, HindIII), see Fig. S8. At larger $v_0$, a modest further reduction of $v_0^{\text{eff}}$ (10–20%) is necessary, possibly reflecting additional contact screening by bulky loop bases. Importantly, no adjustment of $d = T$ was required in this theory-simulation comparison.

### 8 Hi-C data analysis

Contact probability curves, $P(s)$, were computed directly from the raw `.cool` files using the `cooltools` library. The logarithmic derivatives were then obtained as $y = d \log P(s)/d \log s$ and smoothed using a one-dimensional Gaussian filter with $\sigma = 1$.

#### Fitting the Hi-C datasets

The theoretical model was fitted to the experimental log-derivatives using two complementary approaches: (i) matching the position of the minimum $(s_{\min}, y_{\min})$ in the log-derivative, and (ii) fitting the curve around the minimum. Both methods produced consistent results for nearly all datasets. Across 33 datasets analyzed in this study, the inferred loop periodicity from dip matching was $T = 169 \pm 36$ kb, while curve fitting yielded $T = 165 \pm 30$ kb. We summarize the fitted values of the parameters using both approaches for all datasets in STable 1.

In the first approach, the experimental dip coordinates $(s_{\min}, y_{\min})$ were compared against theoretical predictions across a grid of parameters $T$ and $v_0^{\text{eff}}$ and the best-fit pair was found. In the second approach, the full theoretical curve was fit to the experimental profile within the interval 8–40 kb, which provides a robust short-scale representation for most datasets. A comprehensive GitHub repository accompanying this paper contains all analysis scripts and Jupyter notebooks reproducing both fitting procedures for each dataset.

Unless otherwise noted, the optimal parameters reported in the main text were obtained using the curve-fitting approach, while the dip-matching method was used for replicate analyses and cell-cycle (mitotic exit) datasets.

#### Inference stability across Hi-C replicates

To test the robustness of our inference scheme, we applied it to multiple Hi-C replicates within a single biological system. We focused on the lineage of datasets from Bonev *et al.*, which span successive stages of mouse neural differentiation. As shown in Fig. S9, replicate-to-replicate variability is low: the relative error in inferred loop periodicity ($T$) and effective fragment length ($v_0^{\text{eff}}$) remains below 10% in both embryonic stem cells (ES; $T = 190 \pm 16$ kb, $v_0^{\text{eff}} = 5.6 \pm 0.4$ kb) and cortical neurons (CN; $T = 208 \pm 11$ kb, $v_0^{\text{eff}} = 8.7 \pm 0.7$ kb). Importantly, CN cells show a $\sim$20% increase in $T$ relative to ES cells, indicating a lower density of cohesin-mediated loops in differentiated neurons. This trend is consistent with the large-scale chromatin reorganization reported during differentiation, including weakened compartment borders and disruption of the Polycomb network.

Neural progenitor cells (NPCs), representing the intermediate state between ES and CN, behave differently. Although their mean values ($T = 180 \pm 27$ kb; $v_0^{\mathrm{eff}} = 8.0 \pm 1.5$ kb) are comparable to the other cell types, they exhibit markedly greater replicate-to-replicate variability. The relative errors reach 12–15% for $T$ and 17–19% for $v_0^{\mathrm{eff}}$, substantially exceeding those of ES or CN cells (Fig. S9B,E). Moreover, NPCs show stronger differences between *in vivo* and *in vitro* conditions: while CN cells are largely robust, NPCs display a ∼20% reduction in loop periodicity in culture (*in vivo*: $T = 180 \pm 27$ kb; *in vitro*: $T = 147 \pm 19$ kb). We suggest that this heightened variability reflects the transitional nature of NPCs, where chromatin folding is more sensitive to both biological state and experimental context.

We also find systematic differences in $v_0^{\mathrm{eff}}$ between cell types, pointing to physical origins. ES cells possess larger nuclei than CN cells (1.73 $\mu$m versus 1.43 $\mu$m, a 21% increase; the data from Bonev et al, 2017), leading to weaker spatial confinement and hence a smaller effective capture radius. In contrast, the more compact CN nucleus enhances fragment clustering, yielding a larger $v_0^{\mathrm{eff}}$. Consistently, we infer $v_0^{\mathrm{eff}} = 5.6 \pm 0.4$ kb in ES cells versus $8.7 \pm 0.7$ kb in CN cells. These observations underscore a key point: while $T$ reflects the intrinsic structural parameter of loopy chromosomes, $v_0^{\mathrm{eff}}$ encodes protocol- and system-dependent influences such as nuclear geometry and fiber persistence, and is therefore inherently non-universal.

#### Analysis of mitotic exit datasets

To monitor changes in chromatin loop properties during mitotic exit, we analyzed two time-resolved Hi-C datasets: one in HeLa cells from Abramo *et al.* and another in mouse erythroblast line G1E-ER4 (G1E) from Zhang *et al.*. In both experiments, cells were synchronized in prometaphase using nocodazole treatment and subsequently released at $t = 0$. The two systems exhibited markedly different exit kinetics: in HeLa cells, approximately 50% of the population reached early telophase within 2.95 h after release, whereas G1E cells progressed to anaphase/telophase within just 25 min.

In the Zhang dataset, individual time points were annotated directly by cell-cycle stage ("ana/telo", "early G1", "mid G1", "late G1"). In contrast, the Abramo dataset labeled samples by elapsed time after release from prometaphase. To facilitate comparison, we used the DAPI- and tubulin-based cell-cycle classification provided in the Abramo's paper to map time points to corresponding stages: metaphase ($t = 2.2$ h), anaphase ($t = 2.55$ h), and telophase ($t = 2.95$ h). For interphase stages, we grouped the raw time points as follows: early G1 ($t = 3$–$3.5$ h), mid G1 ($t = 4$–$5$ h), and late G1 ($t = 6$–$7$ h). Within each group, replicate curves were averaged to quantify intra-stage variability (see Fig. S10).

We then applied the theoretical expression for the log-derivative of the contact probability (Eq. 4, main text) to each experimental profile to extract the best-fit parameters $T$ (loop period) and $v_0^{\mathrm{eff}}$ (effective fragment length) from the position of the dip (Figs. S10, S11). In both datasets, the inferred loop density exhibited a transient peak pattern, consistent with condensin unloading followed by progressive cohesin reloading. Specifically, in the Abramo dataset, loop density reached its minimum at telophase—coinciding with the transition between condensindominated and cohesin-dominated architectures—mirroring the interpretation of Abramo *et al.* ("intermediate folding state").

In the Zhang dataset, the “ana/telo” time point combines anaphase and telophase, and ChIP–seq profiles for RAD21 showed that cohesin occupancy remains low until early G1 (Fig. 3C in Zhang *et al.* paper). Consistent with this observation, our analysis revealed the largest loop period (corresponding to the lowest loop density) at the “early G1” stage, followed by a progressive decrease in $T$—indicating the gradual buildup of cohesin-mediated loops—through mid and late G1. This temporal trajectory reflects the slower establishment of interphase loop architecture in mESCs, in contrast to the more rapid cohesin reloading observed in HeLa.

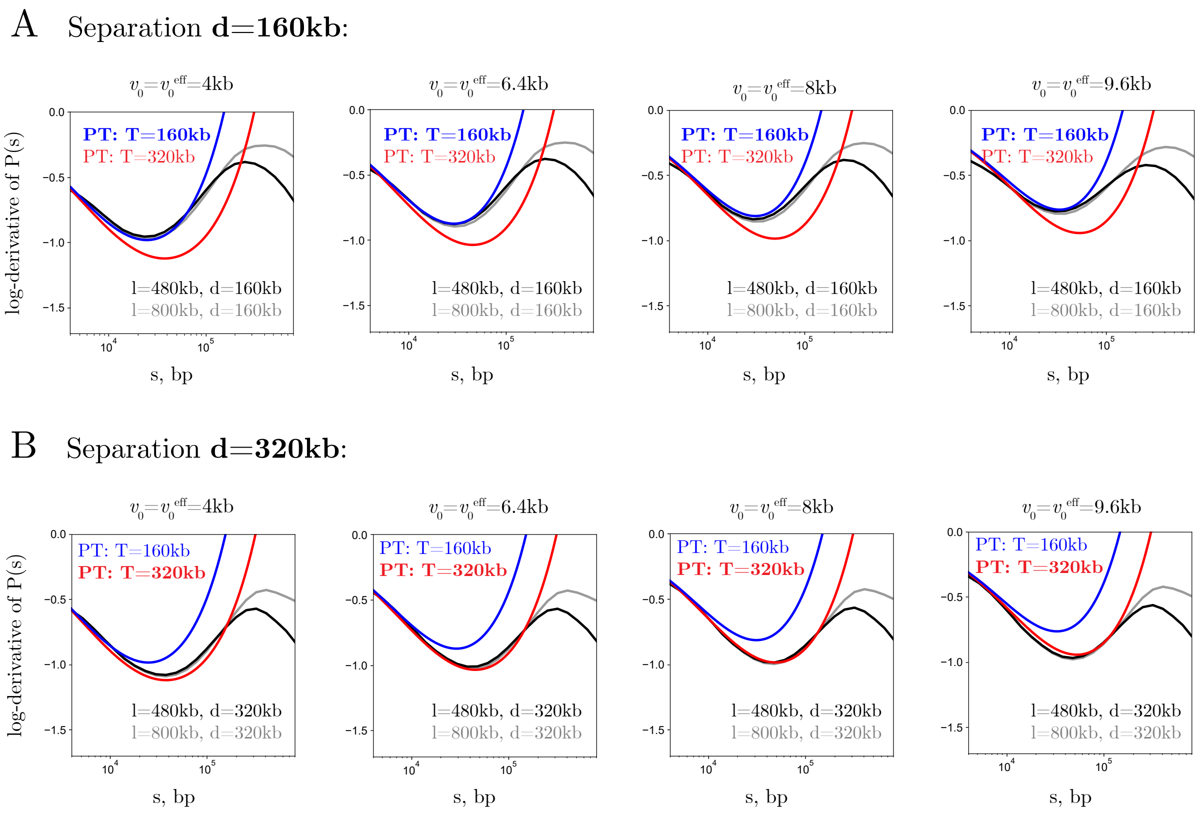

Figure S6: Log-derivatives of contact probabilities $P(s)$ from simulations of ideal (phantom) chains with actively extruded loops. (A) Mean cohesin separation $d = 160$ kb. (B) Mean cohesin separation $d = 320$ kb. In each panel, black/gray curves correspond to simulations with different fragment lengths $v_0$ and cohesin processivities $l$, as indicated. The colored lines (blue and red) represent theoretical predictions with loop period $T$ and effective fragment length $v_0^{\rm eff}$ from perturbation theory (PT). Across both $d$ values, PT accurately reproduces the short-scale behavior and the location of the dip with $T = d$ and $v_0^{\rm eff} = v_0$. Notably, variations in processivity $l$ leave short-scale folding unchanged, affecting only larger genomic separations around the peak.

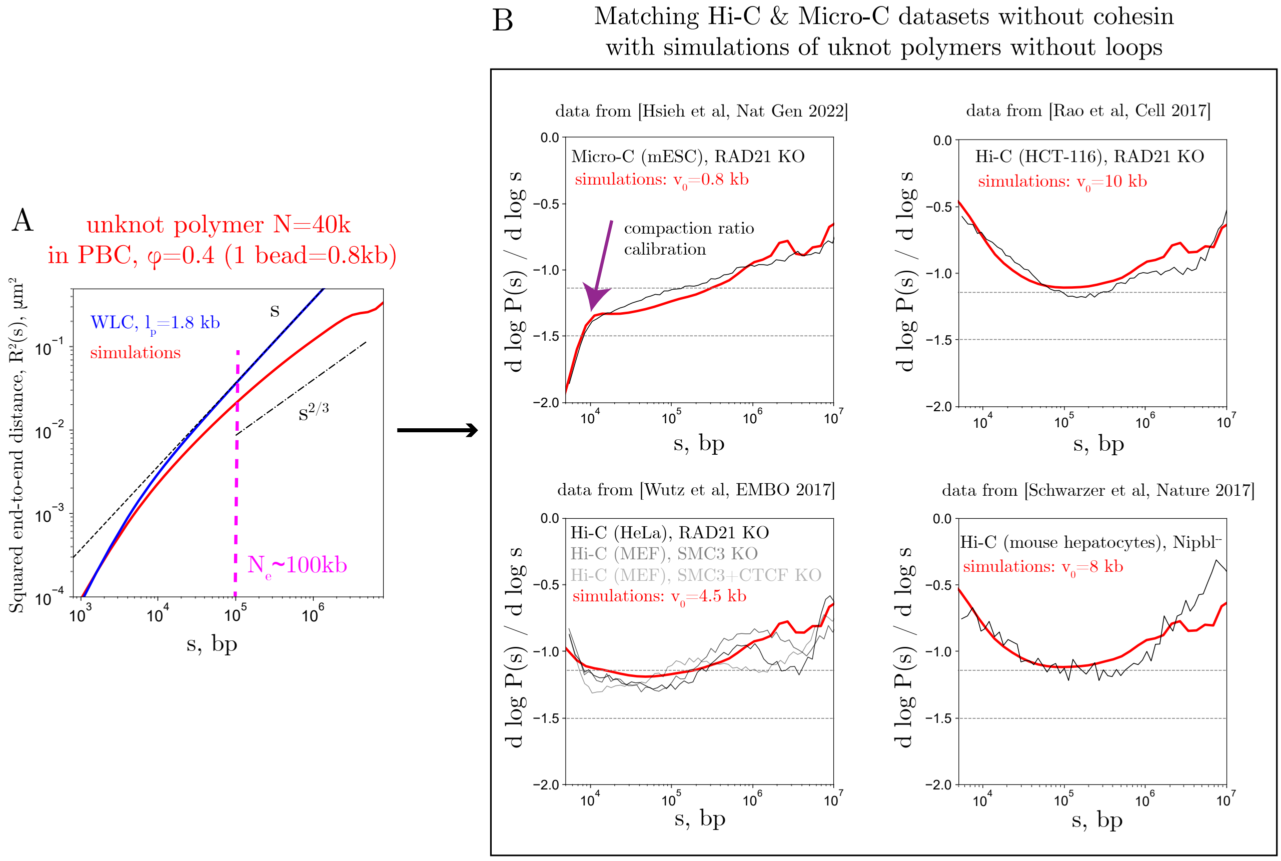

Figure S7: Calibration of crumpled-chain loop-free simulations (unknotted ring polymer, $N = 40{,}000$ beads in periodic boundary conditions) at volume density $\varphi = 0.4$ against Micro-C and Hi-C data with removed cohesin. (A) Squared end-to-end distance $R^2(s)$ of polymer segments as a function of contour length $s$. Simulation results (red) are compared with a worm-like chain (WLC) fit (blue). At short $s$, the WLC fit yields a persistence length of $l_p \approx 2.25$ monomers ($\approx 1.8$ kb using the compaction ratio of 800 bp/monomer). At larger $s$, $R^2(s)$ crosses over to the crumpled-scaling regime $s^{2/3}$, not captured by the WLC, revealing an entanglement length of $N_e \approx 100$ kb. (B) Log-derivatives of contact probability $P(s)$ from simulations (red) matched against Hi-C and Micro-C datasets lacking cohesin, by varying only the fragment length $v_0$. Calibration against Micro-C (top left panel) fixes the bp/monomer conversion and reproduces the characteristic wedge-shaped profile of the experimental curve ($v_0 = 0.8$ kb). Increasing $v_0$ in simulations reproduces cohesin-depleted Hi-C datasets (other panels), demonstrating that variation in $v_0$ alone reconciles cohesin-free Micro-C and Hi-C.

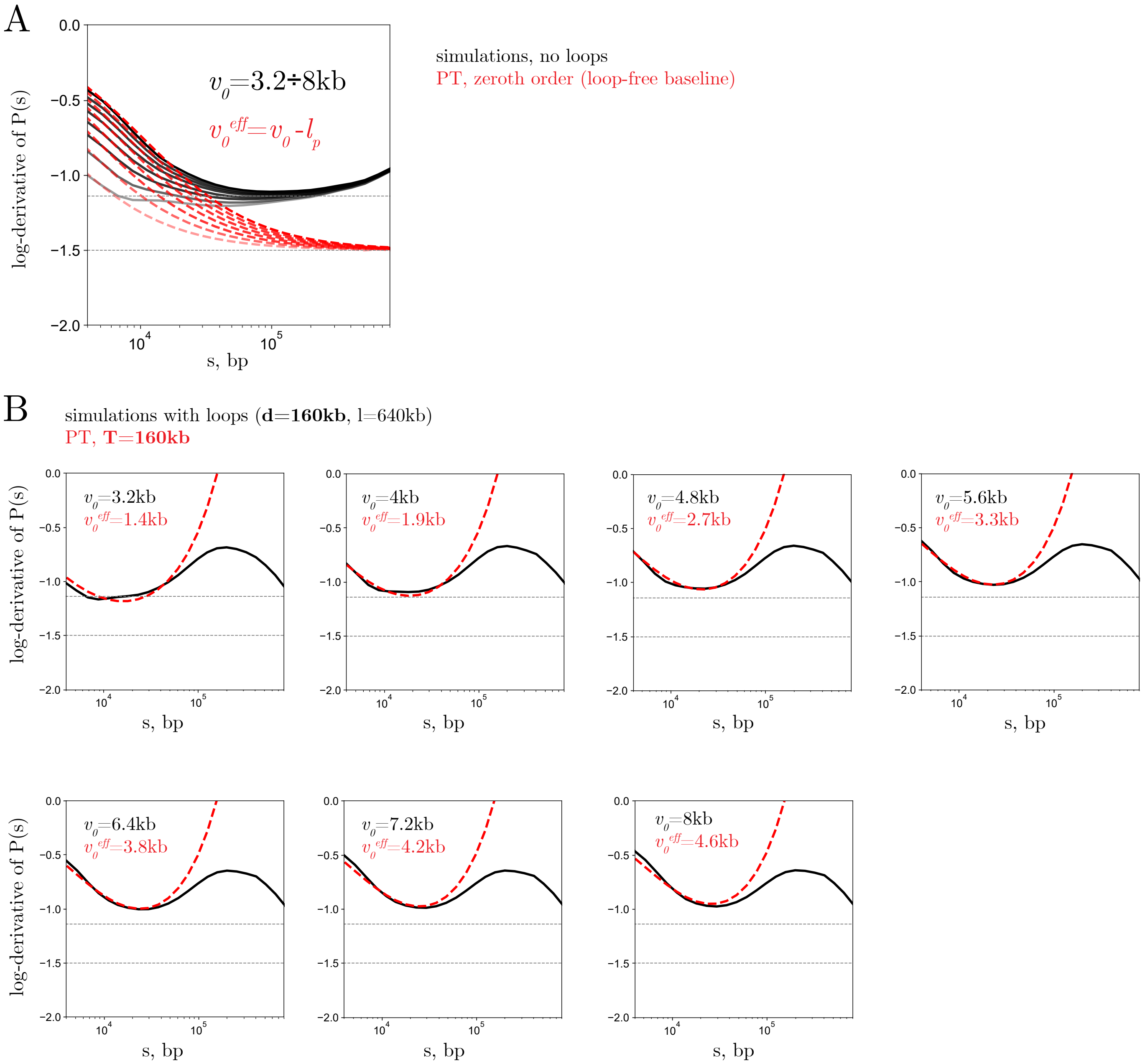

Figure S8: Comparison of perturbation theory (PT) with simulations of crumpled chains. (A) Loops-free case: PT curves, corresponding to the zeroth order (red), compared with simulations of crumpled chains without loops (black). For fragment lengths $v_0 > l_p$, the values of parameter $v_0^{\rm eff}$ from PT closely follow the persistence correction $v_0^{\rm eff} = v_0 - l_p$. (B) Loops-present case: PT curves for $T = 160$ kb and various $v_0^{\rm eff}$ compared with simulations of crumpled chains with actively extruded loops at separation $d = T = 160$ kb and processivity $l = 640$ kb. Across fragment lengths, PT faithfully describes the simulations and reproduces the correct loop density $T^{-1}$. Best-fit $v_0^{\rm eff}$ values are well captured by the persistence correction up to $v_0 \approx 5$–6 kb. For larger fragments, an additional reduction of $v_0^{\rm eff}$ is required. This additional correction leaves the inferred loop density $T^{-1}$ unaffected.

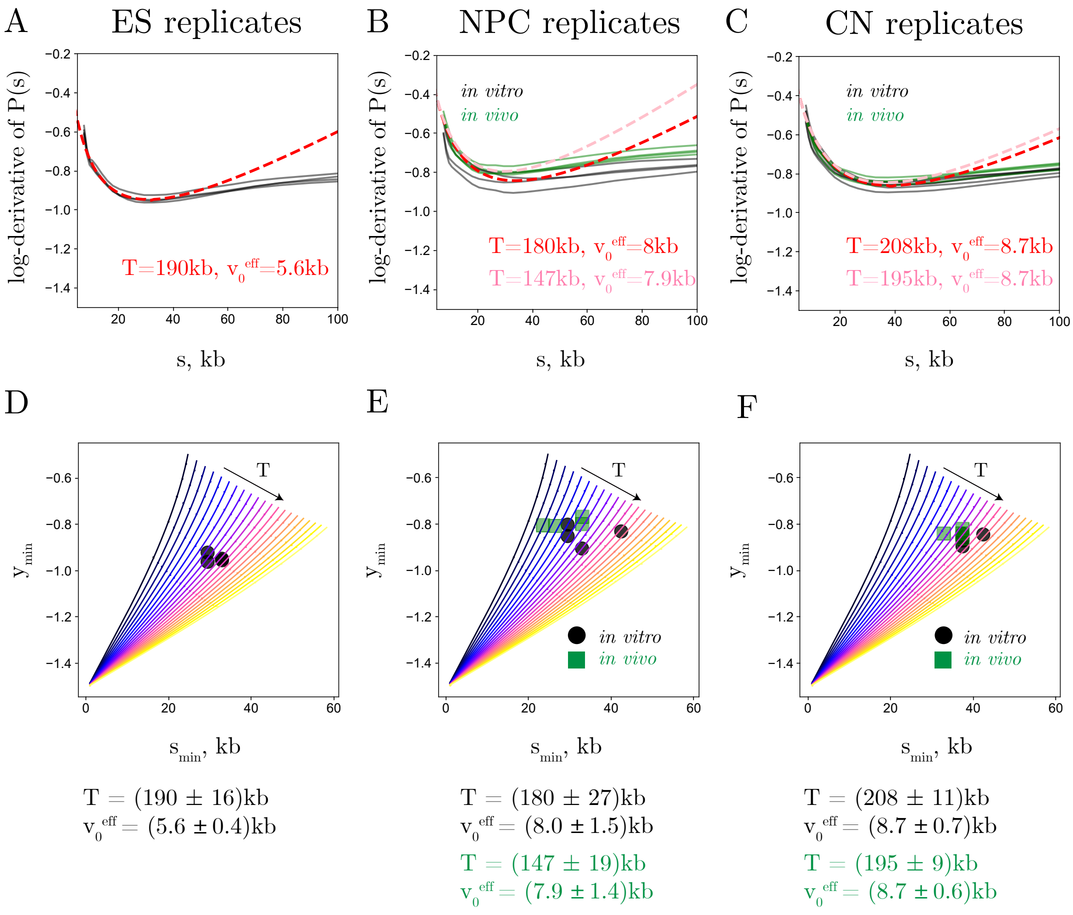

Figure S9: Analysis of Bonev *et al.* dataset. (A–C) Log-derivatives of $P(s)$ for all available replicates of embryonic stem cells (ES), neural progenitor cells (NPC), and cortical neurons (CN). In vitro replicates are shown in gray, in vivo replicates in green. Dashed red curves show perturbation theory (PT) fits to replicate-averaged profiles; fitted parameters are indicated in the legends. (D–F) Positions of experimental minima corresponding to (A–C). From these minima, the loop periodicity $T$ and effective fragment length $v_0^{\text{eff}}$ were inferred for each cell type, yielding robust estimates of cohesin loop density across differentiation stages.

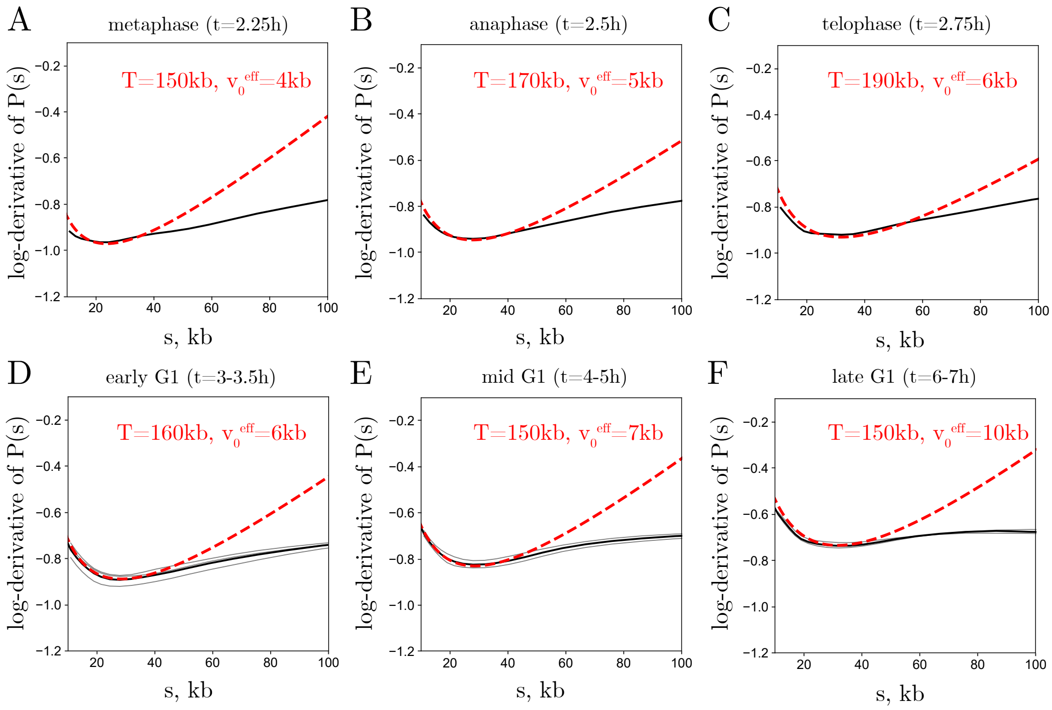

Figure S10: Analysis of Abramo *et al.* dataset. Log-derivatives of $P(s)$ (black) and theory fits (dashed red) for successive stages of mitotic exit. For G1 (D-F), raw time points were grouped into early, mid, and late stages as indicated in the panel titles; theory fits are shown for the group averages to keep track of variations within the temporal groups and the inference stability. Tracking the minima across time enables inference of changing loop periodicity $T$, revealing a transient reduction in loop density during condensin unloading through telophase (t=2.75h), followed by recovery as cohesin loads in G1.

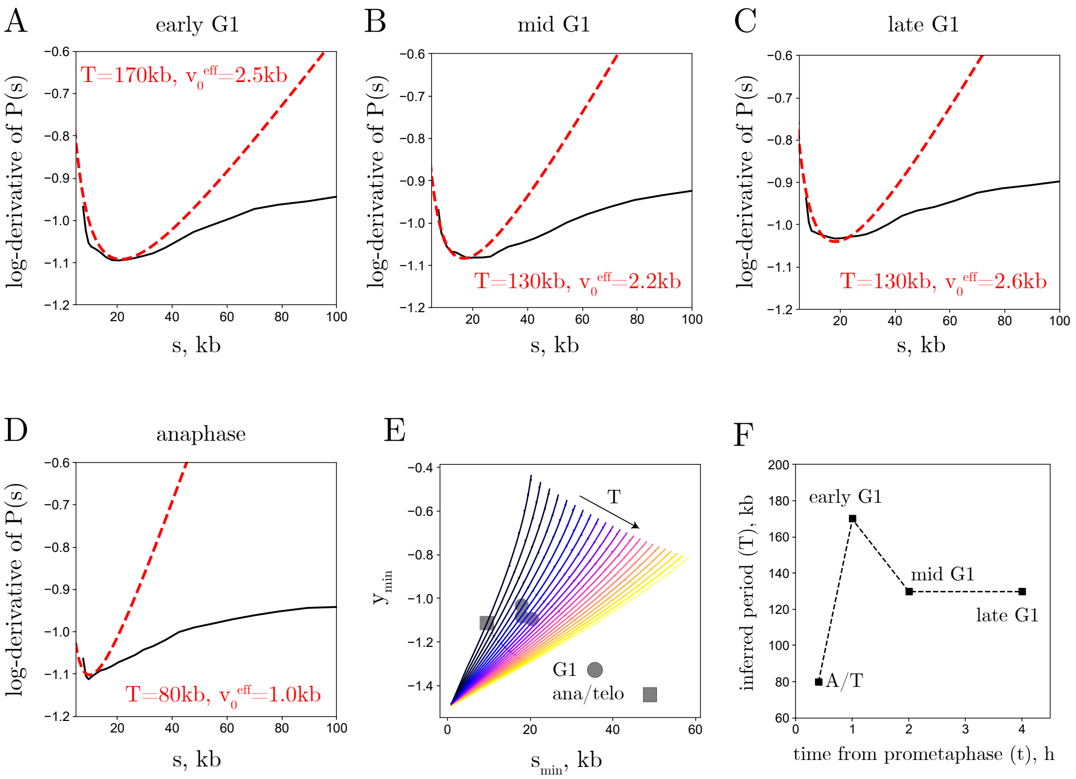

Figure S11: Analysis of Zhang *et al.* dataset. (A–D) Log-derivatives of experimental $P(s)$ (black) with theory fits (dashed red) at successive stages of mitotic exit. (E) Evolution of the minima compared with theory predictions, from which loop periodicity $T$ and loop density $T^{-1}$ were inferred. (F) Schematic of loop-period dynamics during mitotic exit, consistent with a transient reduction of cohesin density followed by stabilization at ∼6 loops/Mb in interphase, analogous to Fig. 6B in the main text.

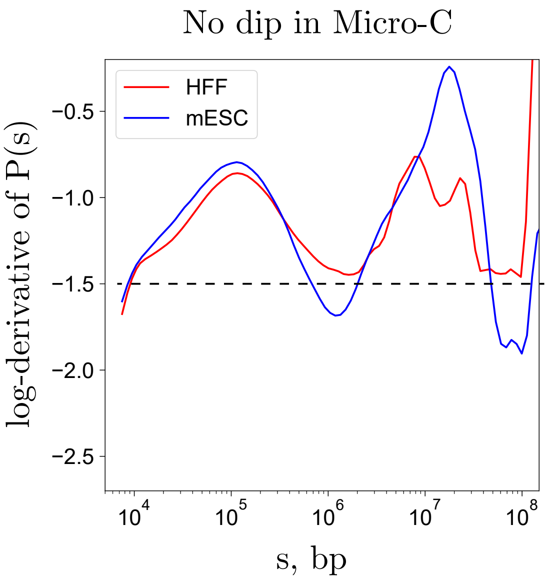

Figure S12: In agreement with the predictions of the theory, in Micro-C data there is no short-scale dip in the log-derivative of $P(s)$: HFF cells (Oksuz et al, Nat. Protocols 2021) and mESC cells (Hsieh et al, Nat. Genetics 2021) are shown. The log-derivative monotonically drops below $-3/2$ at $s < 10$kb, because the restriction fragment in Micro-C is smaller than persistence length, $l_p$.
